# Nervous System and Tissue Polarity Dynamically Adapt to New Morphologies in Planaria

**DOI:** 10.1101/815688

**Authors:** Johanna Bischof, Margot E. Day, Kelsie A. Miller, Joshua LaPalme, Michael Levin

## Abstract

The coordination of tissue-level polarity with organism-level polarity is crucial in development, disease, and regeneration. Exploiting the flexibility of the body plan in regenerating planarians, we used mirror duplication of the primary axis to show how established tissue-level polarity adapts to new organism-level polarity. Tracking of cilia-driven flow to characterize planar cell polarity of the epithelium revealed a remarkable reorientation of tissue polarity in double-headed planarians. This reorientation is driven by signals produced by the intact brain and is not hampered by radiation-induced removal of stem cells. The nervous system itself adapts its polarity to match the new organismal anatomy in these animals as revealed by distinct regenerative outcomes driven by polarized nerve transport. Thus, signals from the central nervous system can dynamically control and re-orient tissue-level polarity to match the organism-level anatomical configuration, illustrating a novel role of the nervous system in the regulation of patterning.

## Introduction

A fundamental question in biology is how spatial patterns are coordinated during dynamic regulation of growth and form. In order to control regeneration, where organisms must repair complex tissues from different starting configurations, it is critical to understand the interaction of patterns across scales, from an individual cell’s cytoskeletal structure to body-wide axial orientation. We used the robust morphogenetic process of regeneration to ask how cell- and tissue-level order is functionally linked to organism-level axial organization. This question is of fundamental biological interest and is also of significant relevance to bioengineering and regenerative medicine, which seek to apply cell-level signals in order to achieve large-scale anatomical outcomes.

Regeneration is a biological process that enables organ remodeling and repair in a number of organisms, most prominent among them planarian flatworms with their ubiquitous ability to regenerate all tissues in their body. While the regenerative process in planarians is well described (Reddien, 2018), we address here two key knowledge gaps. First, it is not known how directional and positional informational cues interact during regeneration – specifically, how tissue-level polarity adapts to changes in organism-level polarity. Second, repatterning and other responses to damage on long timescales have received comparatively little attention; here, we characterize such long-term changes and explore the mechanisms that drive them.

In addition to forming new structures, planarian regeneration requires that the remaining tissues adjusts to their new position in the organism in a process termed repatterning (Adell, Cebria, & Salo, 2010; Morgan, 1901; Reddien & Sanchez Alvarado, 2004). In normal planarian regeneration, this repatterning involves the appropriate scaling of pre-existing organs to the animal’s new size, re-establishment of the correct length/width ratio of the animal, and adaptations of expression patterns of positional control genes (PCGs) as tissues adapt to their new position within the organism (Witchley, Mayer, Wagner, Owen, & Reddien, 2013). The repatterning is more apparent in animals induced to regenerate with atypical morphologies, such as double-headed planarians (Durant et al., 2017; Oviedo et al., 2010) where large sections of tissue in the original posterior of the animal undergo an anteriorization process, including the duplication of structures such as the pharynx.

The flexibility of the planarian body plan enables investigation of the response of existing tissues to new signaling environments via the induction of double-headed animals with mirror-image duplication of the anterior-posterior axis from fully formed single-headed animals, which is not possible in other systems used to study tissue polarity, such as *Drosophila* wings and *Xenopus* embryos (Devenport, 2014; Werner & Mitchell, 2012b). Planar cell polarity (PCP) signaling is one of the highly conserved pathways that underlie tissue polarity in distantly related animals (Davey & Moens, 2017; Devenport, 2014; Marshall & Kintner, 2008; Wallingford, 2012). PCP signaling plays a key role in establishing and maintaining polarity in tissues, for example in the setting of polarity in multiciliated epithelia (Brooks & Wallingford, 2014; Meunier & Azimzadeh, 2016; Spassky & Meunier, 2017) and cancer suppression (Lee & Vasioukhin, 2008). Alternative mechanisms for setting polarity are also widespread, such as the morphogen gradients driving blastoderm polarization in *Drosophila* embryos.

PCP signaling determines the polarity of many tissue structures, among them multiciliated epithelia. Cilia are microtubule-based structures that grow out of basal bodies attached to the cell membrane and which, through a dynein motor driven process perform a beating motion. The orientation in which the basal body is attached determines the beat direction of the cilium (Y.-H. Chien et al., 2013). The establishment of ciliary polarity, via the orientation of the basal bodies, relies on both molecular signals from PCP transmitted via the cytoskeleton (Kunimoto et al., 2012; Wallingford, 2010; Werner et al., 2011; Werner & Mitchell, 2012a), as well as hydrodynamic cues (Y. H. Chien, Srinivasan, Keller, & Kintner, 2018; Mitchell, Jacobs, Li, Chien, & Kintner, 2007). While cilia in single-celled organisms can reverse their beat directions (Iwadate & Suzaki, 2004; Noguchi, Kitani, Ogawa, Inoue, & Kamachi, 2005), there have not been any reports of cilia in multiciliated epithelia cells changing their beat direction after it has been established during development. Ciliary function is thus a sensitive and convenient readout of the underlying epithelial planar organization.

Multiciliated epithelia fulfill many physiological functions (Spassky & Meunier, 2017), including the clearing of mucus in the airway (Konishi et al., 2016), the movement of cerebrospinal fluid in the brain ventricles (Ohata & Alvarez-Buylla, 2016), and motility in the larval forms of many marine animals (Veraszto et al., 2017). In planarians, the multiciliated epithelium on the ventral surface is responsible for the main mode of movement of the worm, allowing smooth gliding (Azimzadeh & Basquin, 2016; Azimzadeh, Wong, Downhour, Sanchez Alvarado, & Marshall, 2012; Rompolas, Azimzadeh, Marshall, & King, 2013; Rompolas, Patel-King, & King, 2010; Rustia, 1925). Classical observational data (Pearl, 1903; Rustia, 1925), as well as modern molecular and theoretical exploration (Vu et al., 2018), showed that in planarians with abnormal morphologies, such as in double-headed or double-tailed animals, cilia polarity is aligned with the polarity of the new body plan via PCP signaling. However, the mechanism by which new morphologies are translated into effects on polarity remains unknown.

The well-described link between PCP signaling and neural development (Goodrich, 2008; Tissir & Goffinet, 2010), and our recent work revealing a dependence on nerve transport for the formation of new body plans in planarians (Pietak, Bischof, LaPalme, Morokuma, & Levin, 2019), led us to investigate how the polarity of the nervous system is affected by atypical body morphologies and how signals from the nervous system impact cilia orientation. Here, we uncover and characterize the remarkable remodeling of the ventral multiciliated epithelium in double-headed planarians, during which cilia beat orientation across large tissue sections is progressively changed, without relying on cell turnover. This reorientation of the epithelial polarity depends on signals from an intact brain. Alongside this repatterning of the multiciliated epithelium, we observed that the overall nervous system polarity is reoriented progressively towards the midpoint of the animal. These results reveal how nervous system structure and tissue planar polarity interact so that pre-existing tissues can be remodeled to accommodate a drastically different body architecture and shed light on the dynamic relationship between directional tissue-level patterning and positional organ pattern on the scale of the entire body.

## Results

### Ciliary beat orientation reveals progressive adaptation to new organismal polarity

Double-headed (DH, mirrored axial polarity) planarians can be created via a variety of methods, including manipulation of bioelectric signaling (Durant et al., 2017; Oviedo et al., 2010) or interference with the Wnt signaling pathway (C. P. Petersen & Reddien, 2008). Newly regenerated DHs (7 days after transection of the single-headed animal) of the species *Dugesia japonica* show developmental differences between their two heads: the head regenerated from the anterior blastema is more fully regenerated than the head positioned on the original tail end of the animal, based on criteria such as eye development, head shape, and blastema coloration (Figure 1A i-iii). We refer to the anterior blastema-derived head as the primary head, while the posterior blastema-derived head is termed the secondary head. The primary head dominates the body movement (Video 1). The morphological differences between the two heads reduce over two to three weeks, eventually leading to both sides being equal in appearance and having equal control over the body movement after about one month (Figure 1Aiv-vi, Video 2). The growth of a secondary head represents an enormous deviation in organismal polarity, and the length of time required to repattern the tissue adjacent to the secondary head provides a unique opportunity to assess the long-term processes that guide remodeling and coordinate axial organ positioning with tissue properties.

**Figure 1:**
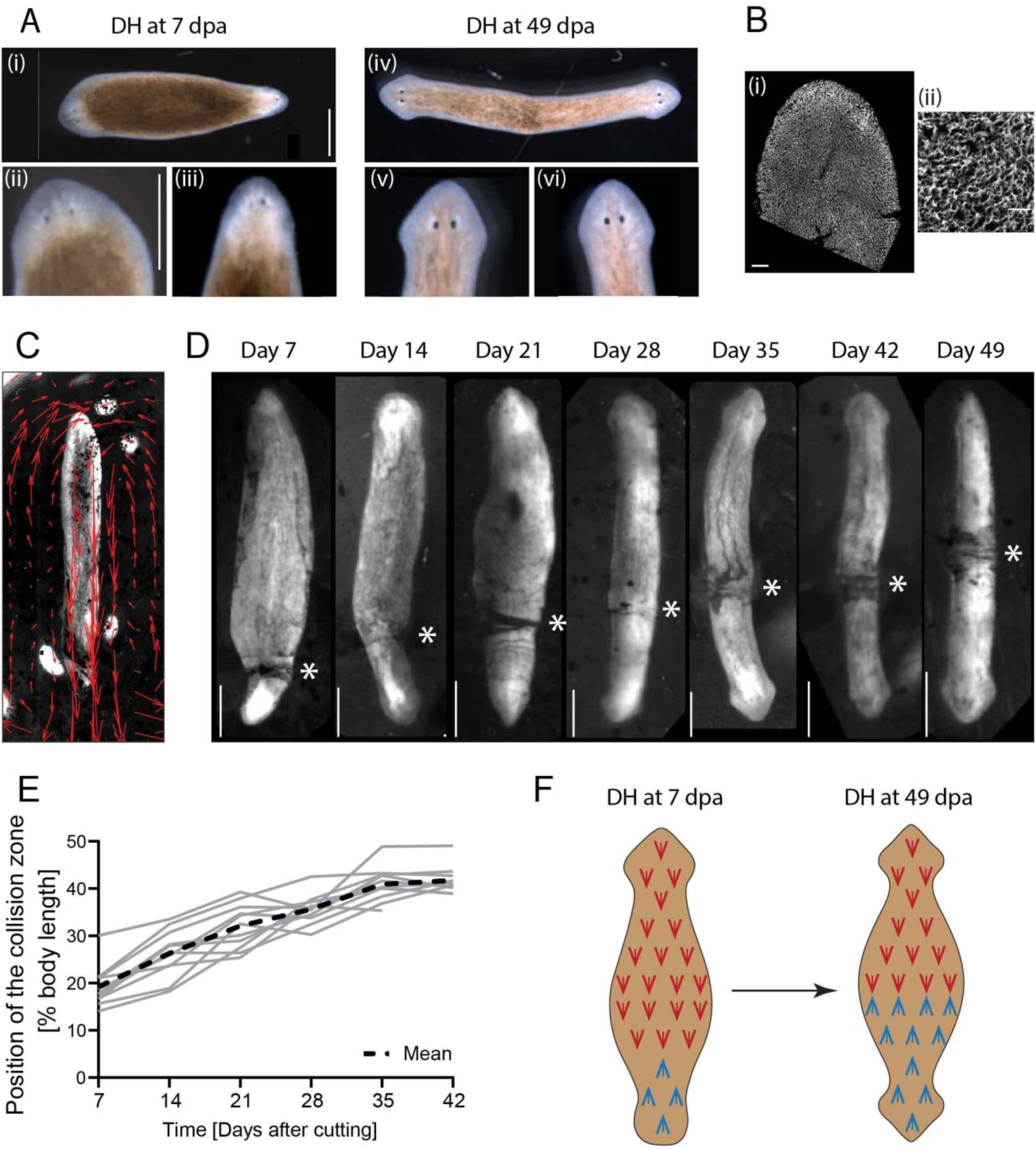
Flow driven by the multiciliated epithelium can be tracked and epithelial polarity dynamically remodels in DH animals. A) (i) Morphological differences in head development in DHs can be observed at Day 7 (days after amputation – dpa), revealing that the primary head (ii) is more developed than the secondary head (iii). (iv) In DHs at Day 49 both heads (v, vi) are equally developed. Scale bars 1 mm. B) Cilia of the multiciliated ventral surface epithelium stained with antibody against acetylated-tubulin at (i) low and (ii) high magnification. Scale bar 100 μm in (i) and 25 μm in (ii). C) Flow driven by cilia beat can be visualized with carmine powder and tracked with particle image velocimetry (PIV) in a single-headed animal (red arrows). D) Opposing flow directions in DH animals lead to the accumulation of particles at the point where the opposing flow fields meet. This “collision zone” (asterisk) repositions towards the middle of the animal from Day 7 until Day 49. E) Plot of position of the collision zone as percentage of total body length for 12 animals individually tracked over time, measured every 7^th^ day, and mean curve (dashed line). N=12, with 5 repeats showing similar pattern. F) Schematic representing the change in cilia orientation in the posterior half of the DH as the cilia orientation changes over time.

The difference in movement patterns between DHs at Day 7 and Day 49 led us to investigate the underlying changes in the multiciliated ventral epithelium, the tissue that drives movement in planarians (Figure 1B) (Azimzadeh et al., 2012; King & Patel-King, 2016; Rompolas et al., 2013). Since cilia beat orientation in multiciliated epithelia serves as a convenient and reliable read-out of underlying tissue polarity (Wallingford, 2010), we adapted an assay to visualize cilia beat orientation in living animals (Pietak et al., 2019; Rustia, 1925). In this assay, worms were placed ventral side up, attached to the underside of the water surface, carmine powder was sprinkled on top of the water, and movement of the powder was observed (Figure 1C, Video 3). Known disruptions of cilia beat function, such as reduction of temperature or 3% ethanol treatment (Stevenson & Beane, 2010), showed immediate cessation or significant reduction of flow, validating our assay (Figure S1). In normal single-headed worms, we observed a complete concordance between the polarity of the epithelium as detected by tracked powder flow and the antero-posterior axial patterning (Figure 1C).

We then tracked the cilia-driven flow in DH animals, which exhibited a clear mirroring of the ciliary polarity, as previously described (Rustia, 1925; Vu et al., 2018). Due to the opposing beat direction in DH animals, the powder used to visualize the flow accumulated at a point along the animal’s body (Figure 1D). We termed this point the “collision zone”, which marks the symmetry point of the ciliated epithelium in the animal. Interestingly we observed that in DHs at 7 days after cutting, the collision zone is located at the base of the secondary head (Figure 1D, Video 4). However, over time, the collision zone progressively shifts along the length of the animal until it reaches the midpoint around 42 days after cutting (Figure 1D, Video 4). This process is highly conserved across all DHs (Figure 1E). The repatterning progresses at a relatively consistent speed from Day 7 to Day 35, before slowing down between Day 35 and Day 42 and stalling following Day 42 (Figure S2A). Repatterning progresses at the same relative speed in animals of different sizes (Figure S2B), suggesting that the repatterning rate is scaling with the size of the animal. Thus, we discovered that the middle of DH worms dynamically adapts its planar polarity over multiple weeks to match the new axial body plan (Figure 1F).

### The signal driving repatterning propagates slowly across the body

The strikingly long repatterning time frame poses the question of whether the repatterning signal is slowly distributed across the animal, or whether the cells of the epidermis require time to adapt to a signal that is present throughout the body at the time of organismal repatterning. To address this, we cut out tissue from the middle of the animal at Day 7 (Figure 2A) to force the formation of new ciliated epidermis in the middle of the body while the body is actively repatterning. If the information is already present throughout the animal, the newly regenerated cells could be expected to regenerate with the correct orientation in place, leading to a faster repatterning or irregular flow pattern in DHs with internal tissue removal. This was not the case. We observed no difference between control animals and DHs with internal tissue removal in both repatterning speed (Figure 2A, Supplementary Table 1) and collision zone/flow pattern (Video 5). This suggests that the signal responsible for the correct pattern is transmitted slowly across the worm rather than epithelial cells slowly adapting to already present information.

**Figure 2:**
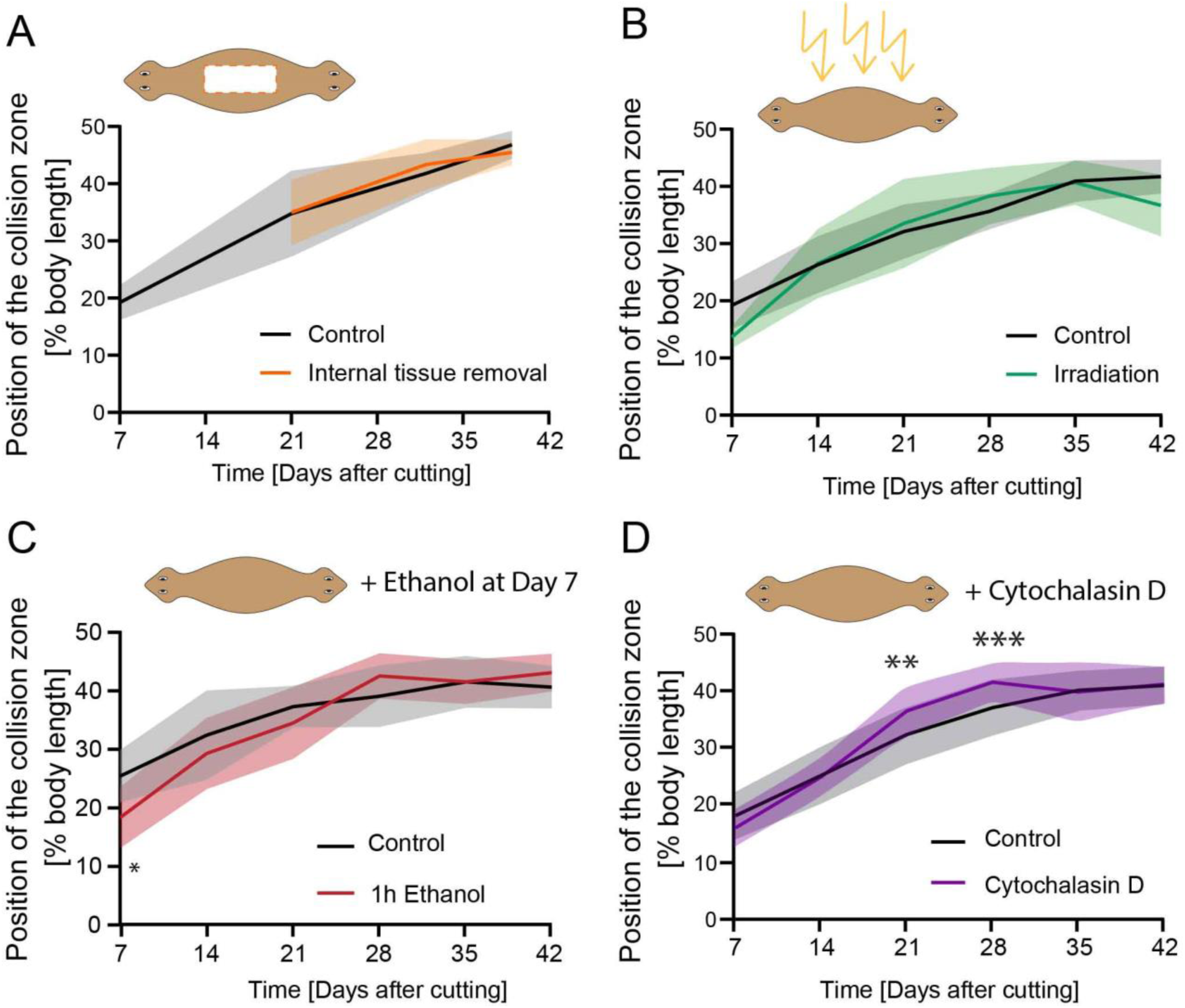
Reorientation of cilia does not rely on cell turnover but is affected by changes to actin cytoskeleton. A) Position of the collision zone over time in animals that had a large portion of tissue removed from their interior (orange line), compared to controls (black); there is no difference in repatterning speed, n=12. B) Repatterning in animals treated with high levels of radiation to prevent new cell formation (green), compared to controls (black); there is no significant difference in repatterning speed, n=12, with 3 repeats showing same pattern. C) Position of the collision zone in DHs which were treated with 3 % ethanol for 1 h at Day 7 to remove external cilia (red); there is no difference in repatterning compared to control (black), except at Day 7 immediately following treatment. n=12. D) Continuous treatment of DH animals with Cytochalasin D (purple) for duration of the experiment compared to control (black) reveals significantly more advanced position of the collision zone for treated animals at Day 21 and Day 28, ** p<0.001, *** p<0.0001. N=12, with 3 repeats showing same pattern. The mean of all samples is plotted. Shaded area represents standard deviation. All experimental worms were compared to age-matched controls.

However, observations in another organ system suggest that this process of slowly transmitted repatterning information across the newly formed double-headed animal is not universal. *De novo* pharynx formation, which is assumed to rely on position control genes (PCG) for its positing (Adler, Seidel, McKinney, & Sanchez Alvarado, 2014), can be observed in DH animals between Day 7 and Day 10. These newly formed pharynges were already positioned correctly in relation to the DH body proportion and orientation. They did not shift their position significantly over time after Day 10 (Figure S3). This suggests that the signals setting the directionality of planar tissue polarity to be in concordance with the new axial morphology may not propagate at the same rate as the signals that determine pharynx positioning.

### Repatterning of the ventral epithelium is not dependent on cell turnover

To investigate whether the repatterning of the multiciliated epithelium requires the replacement of wrongly oriented cells with new, correctly oriented cells, we treated animals 7 days after DH induction (as soon as DHs had completed regeneration) with high doses of radiation (~200 Gy), which is known to kill all stem cells (neoblasts) and thereby prevent all new cell formation in planaria (Wagner, Wang, & Reddien, 2011). Complete removal of neoblasts following irradiation was confirming by absence of staining for actively dividing cells (Figure S4), a lack of regeneration following cutting, and death around 40 days after irradiation. We observed no difference in the repatterning speed of these irradiated animals compared to controls over the entire repatterning time period (Figure 2B, Supplementary Table 2); moreover, irradiated animals had collision zones positioned almost at the midpoint before they died. While some reorientation could be explained by the presence of progenitor cells at the time of irradiation, the full repatterning observed over the long time frame, strongly suggests that cilia reorientation in repatterning DHs happens via the intracellular reorientation of the pre-existing cilia rather than via the replacement of the misoriented cells and is not dependent on neoblast activity.

### Tissue repatterning is actin-dependent

To investigate how the reorientation of cilia occurs in existing cells, we first removed the protrusive components of the cilia using a low dose ethanol treatment (Stevenson & Beane, 2010). DHs were treated at Day 7 for 1 hour with 3% ethanol and did not show different repatterning speeds compared to untreated controls except at Day 7, measured 4 hours after ethanol treatment, indicating potential incomplete regeneration of cilia in this time frame (Figure 2C, Supplementary Table 3). This suggests that repatterning cannot induced by removal and rebuilding of the external cilia structures.

Given the previous discovery that serotonergic neurons are responsible for controlling cilia beating in planarians (Currie & Pearson, 2013; Marz, Seebeck, & Bartscherer, 2013), we explored whether the same neurons also controlled cilia reorientation. We treated DHs with serotonin (Day 7 to Day 14), the serotonergic neurotoxin 5,7-dihydroxytryptamine (5,7-DHT)(Day 7 to Day 42), or the serotonin reuptake inhibitor fluoxetine (Day 7 to Day 42), but observed no difference in repatterning speeds for any of those treatments (Figure S5, Supplementary Table 4–6). Serotonin treatment itself drastically reduced cilia beating (Video 6) but after wash-out there was no difference in the position of the collision zone compared to controls (Figure S5C). These data suggest that cilia repatterning and cilia activity may be controlled via distinct neural circuits, although the sustained nature of our treatment may have differential effects compared to pulsatile signals arising naturally.

To investigate whether a disruption of the cytoskeleton can impact repatterning speeds, we treated repatterning animals with low doses of the highly specific actin depolymerizer Cytochalasin D (Schliwa, 1982) continuously from Day 7 to Day 42 (Figure 2D, Supplementary Table 7). We observed a significantly advanced collision zone at Days 21 and 28 in the Cytochalasin D treated animals (Figure 2D, Day 21 position of collision zone at 37% in Cytochalasin D treated animals and 33% body length in control, Day 28: 42% vs 37% for treated and control animals respectively, Supplementary Table 7). The final endpoint reached was the same for both control and Cytochalasin D treated animals, suggesting that actin depolymerization does not impact the underlying signal and final outcomes, but rather speeds up the process by which the cilia reorientation occurs.

### Repatterning is dependent on the presence of the head

Given the recent characterization of the importance of brain-derived signals in morphogenesis (Herrera-Rincon, Pai, Moran, Lemire, & Levin, 2017), and the observation that cilia repatterning extends from the new head towards the middle of the body (Figure 1D), we hypothesized that one or both brains in a DH animal may be driving remodeling. Firstly, we characterized if the relative smaller size of the secondary brain might be affecting its ability to drive cilia repatterning. We measured brain size in the two brains of DHs using immunostaining for synapsin at different timepoints and quantifying the length of the brain (Figure 3A and B). We observed that early during DH development, between Day 7 and Day 21, there is a great variability in the relative brain lengths between the two heads of the same animal. This indicates that some animals have a much more developed primary brain compared to their secondary brain, while other DH animals have evenly sized brains already at Day 7. The variability reduced progressively from Day 7 to Day 21, when it reached stable values that did not change over the next 22 days until Day 43. We did not observe any correlation between large differences in relative brain size and either positional origin of the fragments cut to induce DHs within the original single-headed (SH) animal (Figure S6A) or overall DH size (Figure S6B). Given the highly consistent repatterning speed across all DHs and the simultaneous diversity of brain sizes, it appears that the secondary brain size does not affect ciliary repatterning.

**Figure 3:**
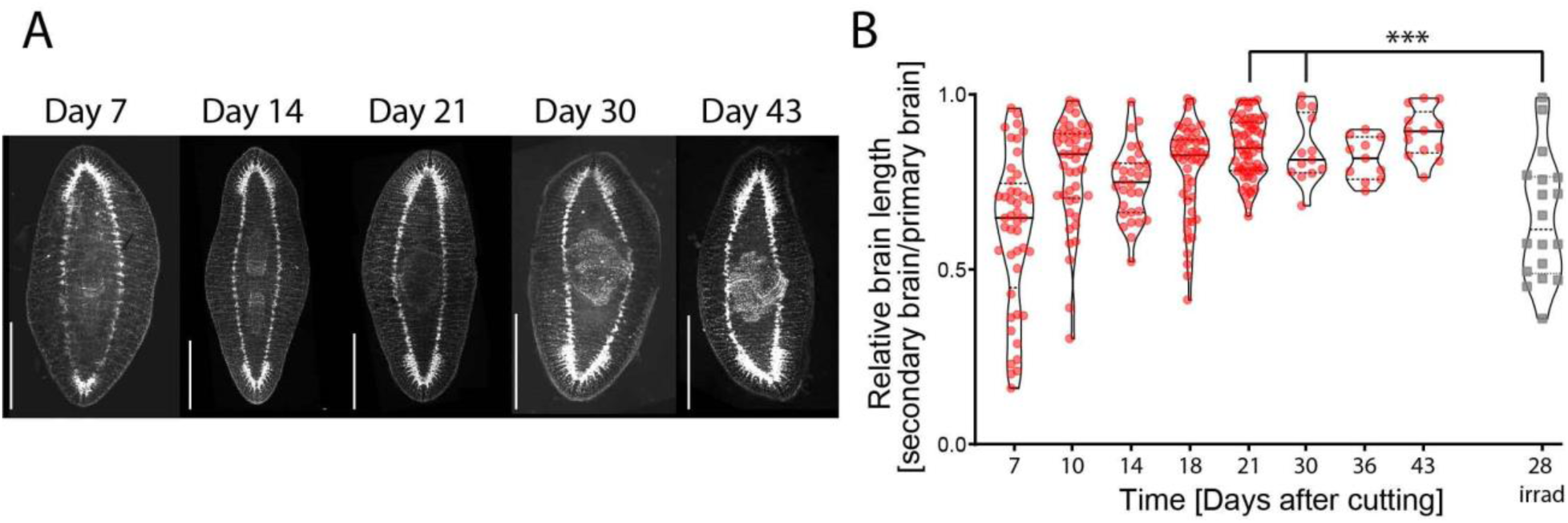
Brain size in DHs changes over time. A) Staining of the nervous system of DH planarians from Day 7 to Day 43 using synapsin antibody staining shows the different sizes of the brain in the primary and secondary head at the early time points. Animals are oriented with the primary head towards the top. B) Quantification of relative brain length in secondary and primary brain over time in untreated double-headed animals (red), and animals irradiated at Day 7 and stained at Day 28 (grey), which shows a reduction in the variability of brain size over time, except for irradiated samples. Brain length was measured in synapsin stains from base of brain where ventral nerve cords (VNCs) widen, to the tip of the brain, where the two halves meet. Scale bar 1 mm. *** p<0.0001

To test the influence of the primary and secondary head on repatterning, DHs at Day 7 were irradiated and then decapitated. Irradiation was performed prior to amputation to prevent regrowth of the removed tissue. As previously shown (Figure 2B), irradiation does not, by itself, impact repatterning. However, removal of the primary head following irradiation led to a significant increase in repatterning speed, with the collision zone being positioned significantly further towards the animal’s midsection compared to controls at all timepoints from Day 14 onward (Figure 4A and C, Video 7, Supplementary Table 8). The removal of the secondary head lead to a reduction of repatterning, with the collision zone advancing significantly less at all timepoints after Day 14 (Figure 4B and C, Video 7, Supplementary Table 9). This indicates that the two heads control cilia reorientation, with the secondary head driving reorientation and the primary head opposing it.

**Figure 4:**
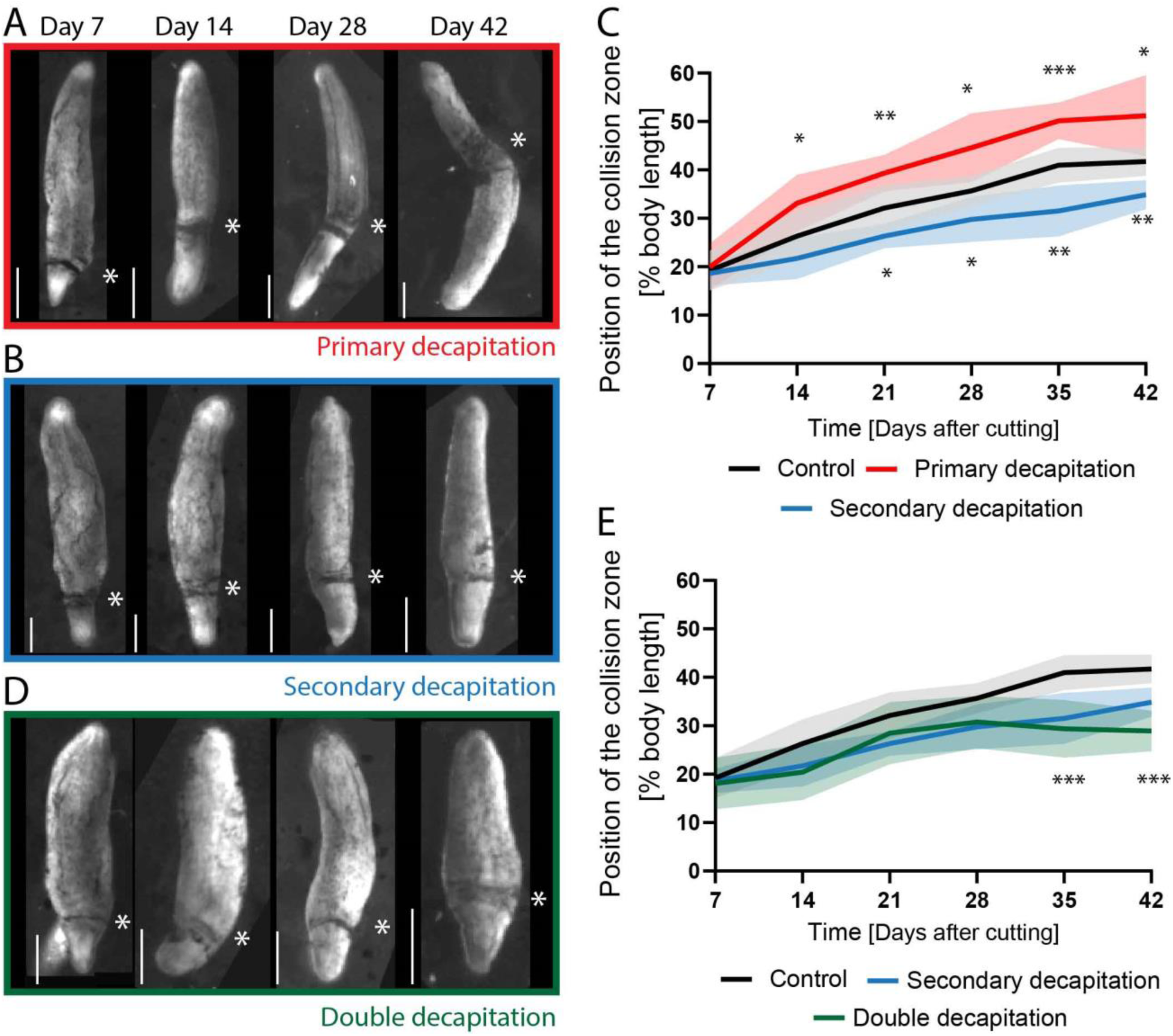
The presence of the heads controls cilia reorientation. A) Position of the collision zone (asterisk) in DH animals irradiated at Day 7 and with their primary head removed at Day 7, 14, 28 and 42, which shows the advance of the collision zone across the midpoint at Day 42. B) Position of the collision zone (asterisk) in DH animals irradiated at Day 7 and with their secondary head removed at same time points. C) Quantification of the position of the collision zone in control animals (black curve) and in animals with either the primary head (red) or the secondary head (blue) removed at Day 7 following irradiation. Removal of the primary head significantly speeds up repatterning, while removal of the secondary head reduces the speed. D) Position of the collision zone (asterisk) in DH animals irradiated at Day 7 and with both heads removed at same timepoints as above. E) Quantification of the position of collision zone in animal with both heads removed (green), compared to control (black) and secondary decapitation (blue). *p<0.01, **p<0.001, *** p<0.0001. N=12, with 2 repeats showing same pattern. The mean of all samples is plotted. Shaded area represents standard deviation. All experiments are paired with their respective controls. Scale bar 1 mm.

To establish whether the reorientation process can take place when both heads are absent, we removed both heads of the DH at Day 7 following irradiation before observing repatterning (Figure 4D and E, Supplementary Table 10). Removal of both heads results in repatterning being significantly slower than in controls, to a similar extent as observed for the removal of only the secondary head. The fact that removal of both heads and removal of only the secondary head give such similar results indicates that the secondary head serves as the main driving force in repatterning – removal of the inhibitory force of the primary brain is not sufficient for repatterning to happen in the absence of the driving force of the secondary brain. At the same time, the collision zone in these double-decapitated animals moves from an average position of 18% body length to 29% body length between Day 7 and Day 42. This may indicate that some head-independent process is able to induce some repatterning or reflect an effect induced before the treatment took effect.

For a limited number of DH animals in which the primary head was removed, we observed that the secondary head dominated reorientation to the point where the collision zone crossed the midpoint of the animal. Irradiated and decapitated animals, however, died before a sufficient number showed repatterning beyond the midpoint to firmly conclude whether removal of the primary head could allow secondary head-mediated repatterning to progress beyond the midpoint far into the primary half of the animal. We therefore took non-irradiated animals and amputated the primary head at Day 7 and then every other day to maintain a worm with no primary head without relying on irradiation. Care was taken to only remove blastema tissue in subsequent amputations. These DHs, which were continuously decapitated on the primary head side, showed repatterning of the cilia leading to the collision zone crossing the midpoint and reaching a collision zone at 86% body length in the most advanced example, while controls did not repattern across the midpoint (Figure S7, Video 8, Supplementary Table 11). Repatterning of cilia reorientation across the whole animal, i.e. collision zone reaching 100%, was not observed, because the animals were not of a sufficient size to allow the assay to be continued after Day 63. The steady slope of the repatterning curve in the continuously repatterning animals however suggests that full reversal of cilia polarity could be achieved. This indicated that there is no set midpoint marked independent of head presence.

The importance of the head in controlling cilia reorientation led us to first investigate whether removal of the anterior pole of the animal, known to express the gene *notum* and to be crucial for tissue organization in planarians (C.P. Petersen & Reddien, 2011), was responsible for the remodeling effect we observed following head removal. We found that removal of just the tip of the secondary head in a non-regenerative irradiated animal did not impact repatterning (Figure S8, Supplementary Table 12). This strongly implicates that *notum* signaling is not required for repatterning of the ciliated epithelium and implicates the brain as the driving factor of repatterning.

It is clear that the presence of the secondary head is crucial in driving the repatterning in the developing DH. To determine whether this is only true during the initial repatterning or whether the presence of the head is required for maintenance of the midpoint, we performed decapitations following irradiation in DHs at Day 49. In these animals the collision zone had reached the midpoint before they were decapitated. In the 21 days following decapitation at Day 49, we did not observe any change in the position of the collision zone (Figure S9, Video 9, Supplementary Table 13). Similarly, decapitating irradiated SH animals did not lead to a change in flow pattern (Video 10). These data indicate that once the position of the collision zone is established, the presence of the brain is no longer required for maintaining its position or organizing cilia beat pattern. Overall, it appears that each brain provides an influence during early establishment of cilia orientation, but once patterning is complete it is not required to maintain the cilia orientation.

### An intact brain is required for driving repatterning

We then investigated whether the overall mass of the brain was a key feature in the strength of this repatterning effect. We removed half of the secondary head laterally (Figure 5A) following irradiation at Day 7. These animals with only half the head removed repatterned at the same speed as animals that had the complete secondary head removed (Figure 5A, Supplementary Table 14). Notably, we never observed an angled collision zone, which indicates that the two lateral halves of the worm are not regulated independently from each other by the brain hemispheres (Figure S10). We also removed half of the head perpendicular to the head-head axis (cut at the eye plane, Figure 5B), which lead to an equivalent reduction in repatterning as lateral half-head removal (Figure 5B, Supplementary Table 15). These data indicate that presence of half the head is not sufficient to complete the function necessary to drive repatterning, independent of which half of the head is removed.

**Figure 5:**
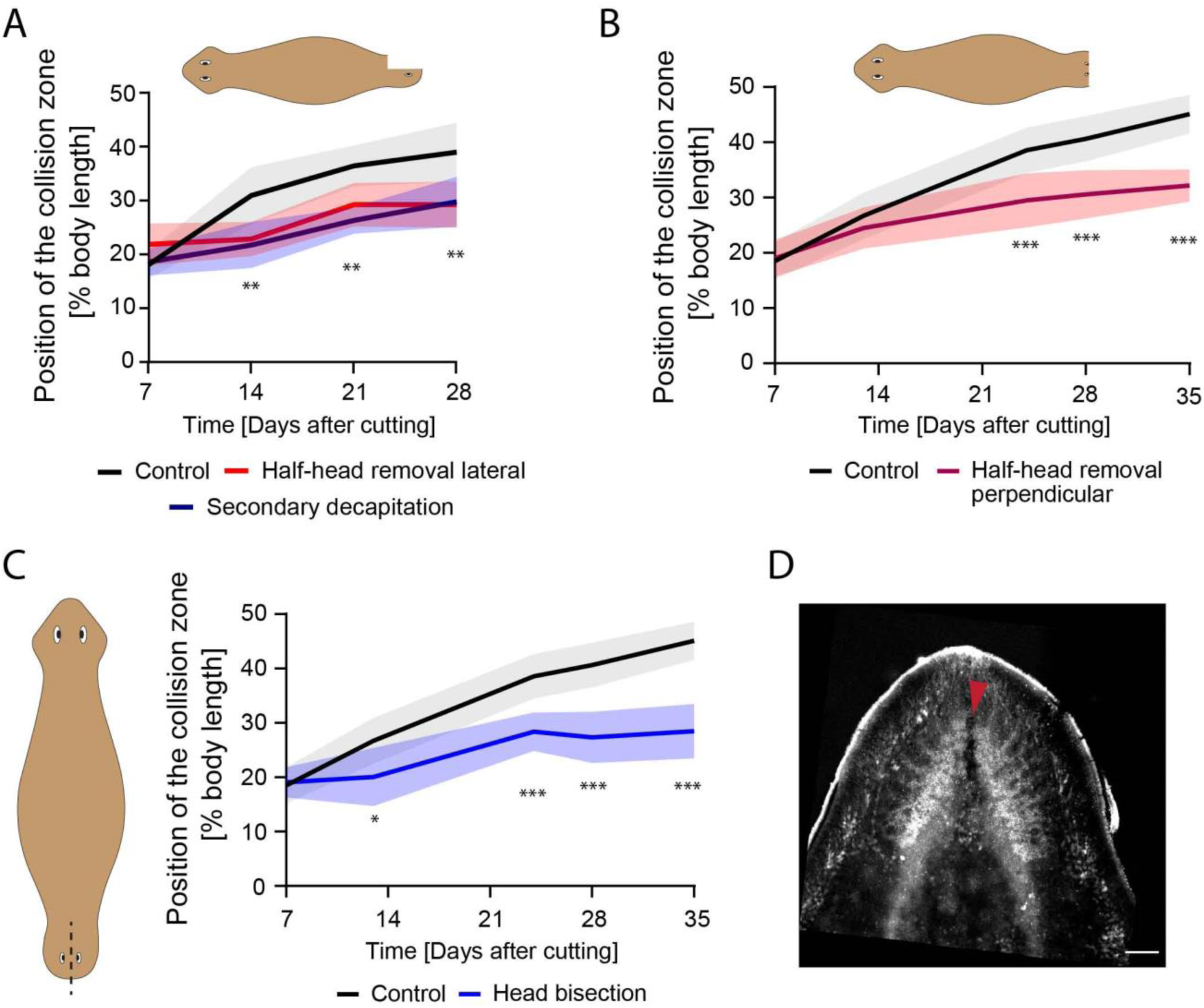
An intact brain is critical for driving repatterning. A) DHs with the lateral half of the secondary head removed following irradiation at Day 7 (red) showed significantly reduced repatterning speed compared to control (black) to the same extent as animals with the entire secondary head removed (blue). B) DHs with the frontal half of the secondary head removed following irradiation at Day 7 (red) showed significantly reduced repatterning speed compared to control (black). C) DHs with secondary head bisected laterally between the eyes without tissue removal following irradiation at Day 7 (blue) repattern significantly slower compared to controls (black). D) Synapsin stain of brain following head bisection without tissue removal. Red arrow highlighting disruption of the brain due to the head bisecting cut. * p<0.01, ** p<0.001, *** p<0.0001. N=12. Plotted is mean of all samples, with standard deviation as shaded area for all experiments with their age-matched controls. Scale bar 100 μm.

The observation that half-head removal was as effective at reducing repatterning as full head removal, led us to ask whether the small size of the secondary brain at the beginning of the repatterning process had an impact on repatterning speed such that a smaller brain cannot send the same signals as a fully formed brain to drive the repatterning. To answer this question, we investigated the brain size of DHs at Day 28 that had been irradiated at Day 7. Irradiation in these animals stopped their brain development and maintained the diversity of brain sizes observed at Day 7 until Day 28, making them significantly different from non-irradiated animals at Day 21 and Day 30 (Figure 3B, Supplementary Table 16). Since irradiated animals repattern at the same rate as controls (Figure 2B), the smaller brain size of the secondary brain is not responsible for slower repatterning.

Given that small intact brains are capable of driving repatterning, while removal of half the brain significantly slowed repatterning, we asked whether any injury to the brain was sufficient to reduce repatterning by bisecting the two halves of the brain in the secondary head via a cut made from the head tip between the eyes in irradiated animals at Day 7 (Figure 5C, D). This cut persisted due to the inability to replace cells following irradiation and was sufficient to significantly reduce the repatterning speed similar as removal of half or the entire head (Figure 5C, Supplementary Table 17). Combined with the previous result, this suggests that it is the intact state of the brain, rather than its overall size, that matters for successful control of cilia reorientation, and suggest that functional signaling across the entire brain may be required for repatterning, as the amount of remaining tissue and any possible diffusion pathways are not affected by simple cut injuries.

### The signals that determine cilia repatterning are transmitted along axial structures

Given the important instructive role of the intact brain in cilia repatterning in developing DHs, we sought to understand by which pathway the repatterning signal is sent from the brain to the rest of the body. To test whether the signal diffused through the body or was transmitted along axial structures, we irradiated DHs at Day 7 and then cut them below either the primary or the secondary head (Figure 6 Ai, Bi and Ci). The wounds healed, reforming the majority of the physical connection between head and body (Figure 6 Aii, Bii, Cii) and allowing for diffusion to take place normally. Using synapsin staining, we showed that following these cuts the ventral nerve cords (VNCs) were severed and the connections did not reform in any of the tested animals due to irradiation-driven lack of new cell formation (Figure 6Aiv, v compared to Biv, v and Civ, v and Figure S11). External wound healing suggests that epidermal connections are reformed, but we did not explore whether muscle connections are reestablished. We found that this axial cut at either the primary or secondary head phenocopied the effect of entire head removal (Figure 6D, Video 11). Compared to controls (Figure 6A), animals with a lateral cut behind the primary head (Figure 6B) reoriented their cilia significantly faster at all timepoints following Day 14 compared to controls (Figure 6D, Supplementary Table 18). At the same time, DHs in which the body was cut behind the secondary head (Figure 6C) repatterned significantly slower, with the collision zone significantly less advanced at all timepoints following Day 14 (Figure 6D, Supplementary Table 19). These data support the hypothesis that the factors driving the repatterning are not diffusing through the mesenchyme but are transported along axial structures disrupted by the cut, such as the VNCs.

**Figure 6:**
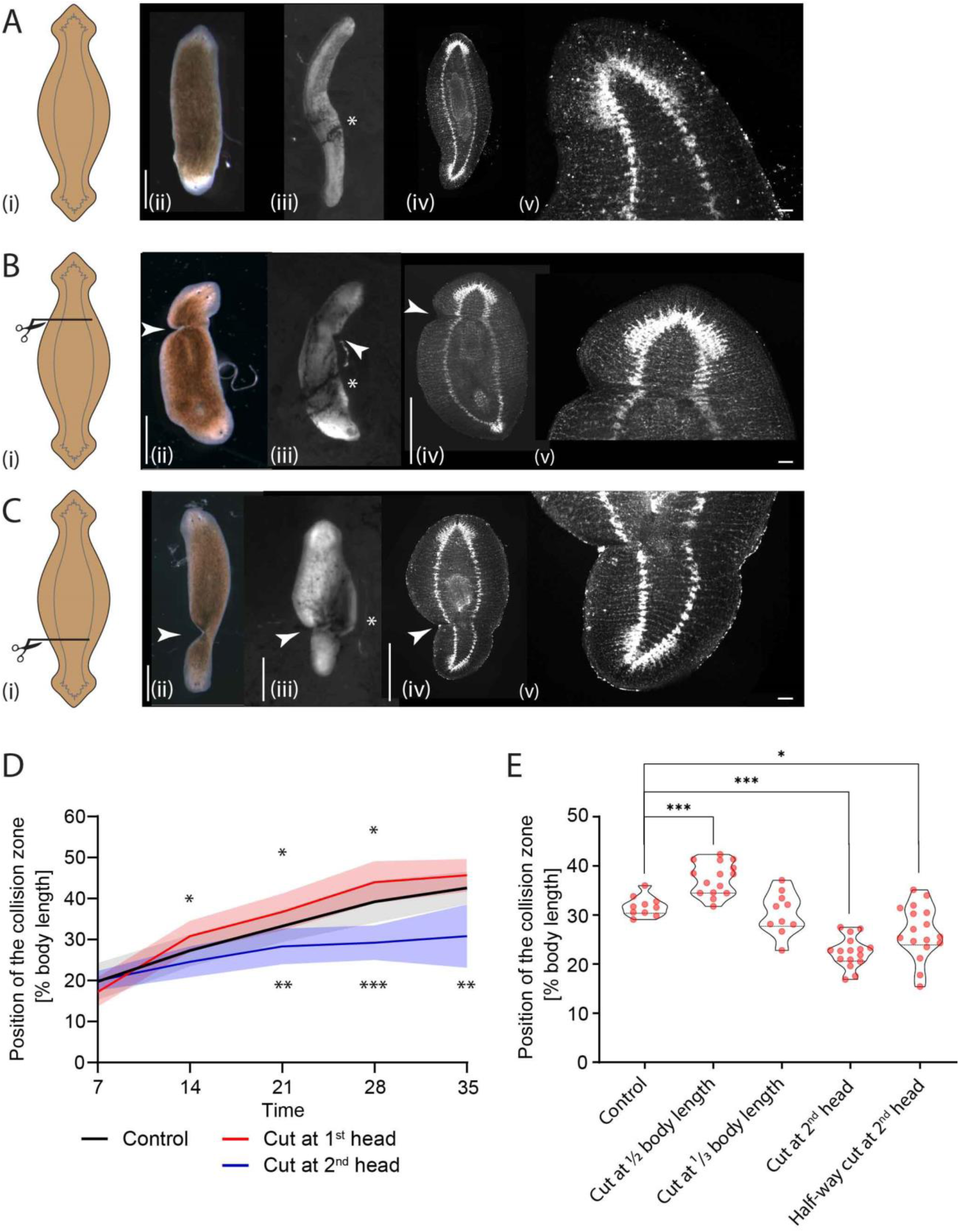
Remodeling-inducing signals from the brain are transmitted across the body. A) Uncut control animals: sketch of animal with CNS (grey) (i), in brightfield at Day 7 (ii), flow assay result of the same animal at Day 28 (iii), synapsin stain in the same animal at Day 28 (iv), and higher magnification of intact VNCs (v). B) DH with cut at primary head: sketch showing cut position with CNS (grey) (i), in brightfield at Day 7 (ii), flow assay result of the same animal at Day 28 (iii), synapsin stain in the same animal at Day 28 (iv), and higher magnification of cut VNCs confirming severance (v). C) DH with cut at secondary head: sketch of cut position with CNS (grey) (i), in brightfield at Day 7 (ii), flow assay result of the same animal at Day 28 (iii), matching synapsin stain at Day 28 (iv), and higher magnification of cut VNCs confirming severance (v). D) Quantification of position of the collision zone in animals with VNCs cut at either the primary head (red) or the secondary head (blue) at Day 7 following irradiation, showing significant difference compared to control (black). N=12. Plotted is mean of all samples, with standard deviation as shaded area, all experiments with their age-matched controls. E) Position of collision zone at Day 21 in animals treated at Day 7 compared to controls, with animals with cut at midpoint, cut at 1/4^th^ body length, cut at base of 2^nd^ head, as well as animals with cut to midline at base of 2^nd^ head – all in the secondary half. *p<0.01, ** p<0.001, ***p<0.0001. Arrowheads mark the cut sites. Scale bar 1mm, in insets 100 μm.

To further confirm that the factors driving repatterning travel the length of the body in long axial structures and to ascertain the spatial dynamics of this influence, we performed lateral cuts at different positions along the head-head axis following irradiation at Day 7 and observed the position of the collision zone at Day 21 (Figure 6E, Supplementary Table 20). When the DHs were cut in the middle of the body, the collision zone was positioned significantly further towards the midpoint compared to controls, mirroring cuts at the base of the primary head, consistent with the signal from the primary head having to be transmitted into the secondary half of the animal to block repatterning. Cuts at 1/3^rd^ body length did not significantly differ from controls (Figure 6E, Supplementary Table 20), likely because the cut was placed close to the position of the collision zone, while animals with cuts between the secondary head and the collision zone (cut at the base of the secondary head) had a significantly lower position of the collision zone (Figure 6E, Supplementary Table 20).

We also performed lateral cuts which only went to the midline, disrupting only one of the VNCs at the base of the secondary head in irradiated worms. In these DHs we found that the depth of the cut did not impact its effect, with a cut severing most of the body resulting in the same repatterning delay as a cut only to the midline (Figure 6E). This result implicates the VNCs as the axial structures required for transmission of the molecules leading to repatterning, as disruption of one VNC is likely to disrupt nerve signaling, consistent with our observations following half-brain removal (Figure 5). Taken together, these data suggest that repatterning relies on the presence of an intact nervous system, potentially for transport of signals that direct repatterning of the cilia in the surrounding area.

### Nervous system in double-headed worms repatterns over time

Given the importance of the nervous system in the adaptation of the tissue polarity of the ventral epithelium, we next explored whether the polarity of the nervous system itself is affected in these repatterning DHs. Our previous work showed that morphogen transport in the nervous system plays an important role in determining regenerative outcomes and that mature DHs have a morphogen transport field that is mirror-symmetrical along the midline (Pietak et al., 2019). In the previous work, we showed that in mature DHs different cutting planes led to distinct regenerative outcomes based on whether the fragment included the midpoint of the animal, where we suggest the two halves of the nervous system meet. Fragments containing the midpoints always regenerate as DHs, while any non-midpoint fragments regenerated as single-headed animals (SHs) (Pietak et al., 2019). Given that the symmetry point of the multiciliated epithelium shifts between immature and mature DHs, we asked whether the symmetry point of the nervous system is similarly affected by assaying its ability to instruct polarity during regeneration.

Our nerve-transport data suggest that if repatterning of the nervous system takes place, the regenerative outcomes of identical amputations performed in 7-day old DHs and 49-day old DHs will be different as the underlying transport field is distinct (Figure 7A). We performed amputations at three different planes: a narrow decapitation directly at the base of one of the heads (around 1/6^th^ of the body length), a cut positioned 1/4^th^ of the way along the head-head axis, and a cut bisecting the DH animals in half. While the decapitations lead to the same regenerative outcomes in both mature and immature DHs, cuts positioned at 1/4^th^ of the body length and at the middle of the body lead to distinct regenerative outcomes for immature and mature DHs.

**Figure 7:**
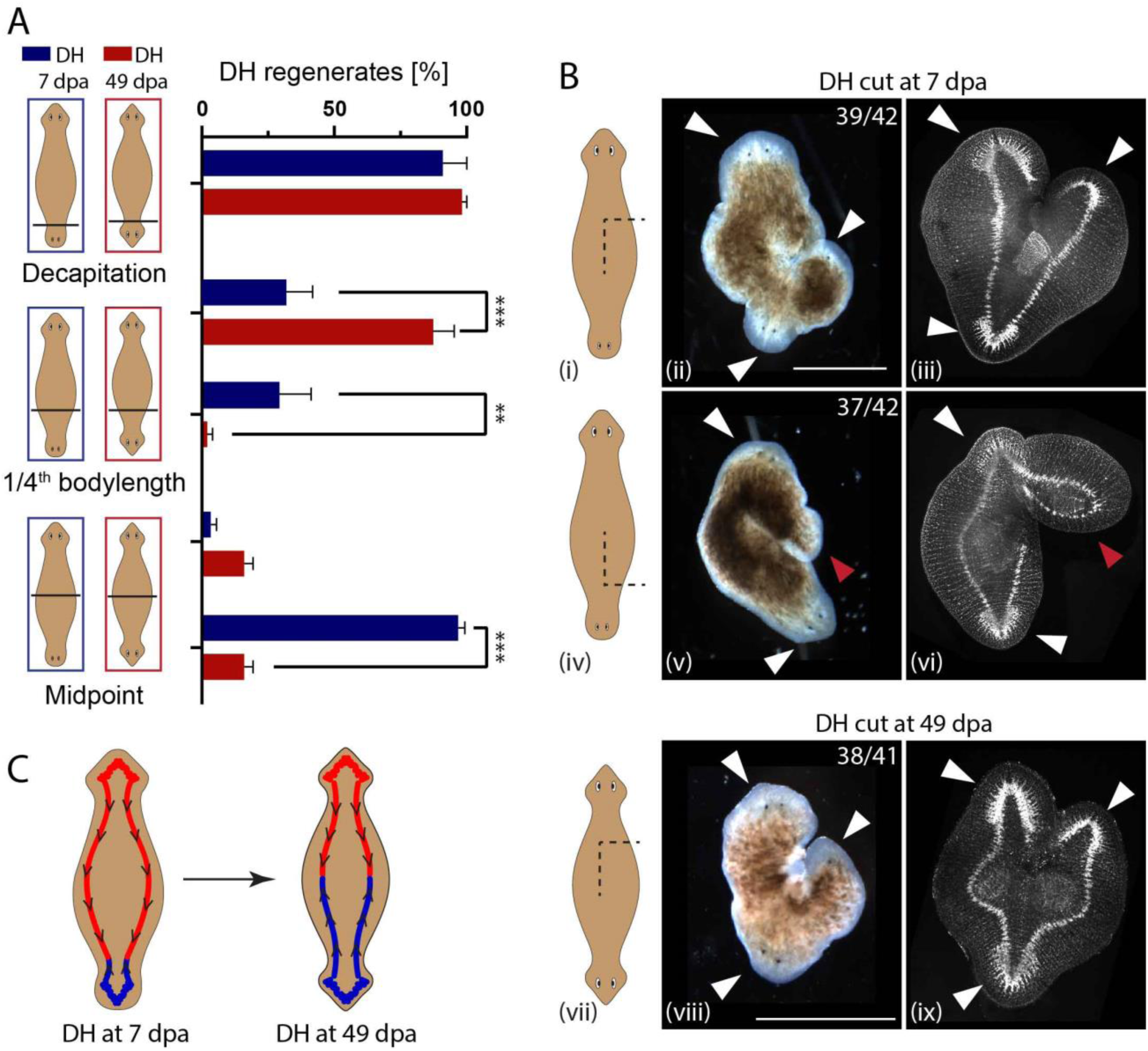
The CNS’ influence over polarity during regeneration changes spatially over time. A) Percentage of DH regenerates from DHs cut at various locations along the body axis. Cuts were performed either in animals at Day 7 (blue) or Day 49 (red) (days after amputation – dpa). Decapitation resulted in 96% and 98% DH for Day 7 and Day 49 animals, respectively. Fragment consisting of 3/4^th^ of the body regenerated at 28% vs. 87% DHs, while the 1/4^th^ fragments regenerated only 29% and 2% DHs for Day 7 and Day 49, respectively. Midway cuts gave 16% DH regenerates for both halves of DHs at Day 49, while DHs at Day 7 had 3% of the primary half regenerating as DHs, while 97% of the secondary halves regenerates as DHs. N=4, with 15 worms per replicate. B) Regenerative outcomes of induced side-outgrowths facing either (i-iii) towards the primary head or (iv-vi) towards the secondary head in DHs at Day 7 and (vii-ix) in DHs at Day 49, shown as sketch (left column), in brightfield (middle column) and in synapsin antibody stain, visualizing the nervous system (right column). Frequency of the observed outcomes is given in each panel. White arrow – head regeneration, red arrow – tail regeneration. C) Model of the nervous system polarity in double-headed animals at Day 7 and Day 49, representing the change in orientation in the secondary half of the animal as it adapts to the new morphology. ** p<0.01, *** p<0.0001, plotted are mean values with error bars representing standard deviation. Scale bars 1 mm.

For the cuts positioned at 1/4^th^ of the body length in mature DHs the larger fragment (containing the midpoint) regenerated significantly more often as a DH compared to immature DHs (87% vs. 31%, p<0.0001, Supplementary Table 21). In immature DHs, both the 1/4^th^ and the 3/4^th^ fragments regenerated as DHs at similar percentages (29% vs. 31%). This suggests that when a cut is placed at around 1/4^th^ of the way from the secondary to the primary head, in approximately one third of the cases the symmetry point of the nervous system is destroyed via cutting, leading to just SH regeneration, while in the other two-thirds of cases the symmetry point is located in either one of the two fragments leading to a regenerative outcome of one DH and one SH, suggesting that the location of the symmetry point of the nervous system at Day 7 is located around 1/4^th^ of the body length away from the secondary head. When mature DHs were cut in half, both halves regenerated mostly as SHs (84% SHs), while for immature DHs the two halves behaved very differently. The half containing the primary head almost always regenerated as SH (96%), while the half containing the secondary head almost always regenerated as a DH (97%) (Figure 7A). These data reveal that the symmetry point of the DH nervous system is positioned very off-center in an immature DH, somewhere around the 1/4^th^ body length point, while it is positioned in the middle in mature DHs, supporting the hypothesis that the nervous system polarity adapts in parallel with the polarity of the ciliated epithelium. The position of the symmetry point of the nervous system in DHs at Day 7 suggested by these experiments at around 1/4^th^ of the body length is consistent with the cilia-flow collision zone located at 27% of body length on average at Day 7.

To show that the underlying orientation of the VNCs in immature and mature DHs is different, we generated side-wounds innervated with a single VNC in both mature and immature DHs (Figure 7B). In the immature DHs, where a distinction between the primary and secondary head could be identified visually, the wounds were induced either facing towards the primary or the secondary head, at about 1/3^rd^ of the body length away from the head they were facing towards. The induced outgrowths were scored for either head or tail identity using both appearance and synapsin staining to visualize the nervous system structures. In immature DHs, side outgrowths facing towards the primary head regenerated with a head 93% of the time (39/42, Figure 7 Bi-iii), while the outgrowths facing towards the secondary head regenerated as tails 88% of the time (37/42, Figure 7 Biv-vi). In mature DHs, no distinction between primary and secondary head remains apparent to guide the positioning of cuts, therefore the comparative outcome to the immature DH result would be 50% heads and 50% tail regenerates in the side outgrowths, as samples are an equal mix of cuts facing the primary and the secondary head. However, we observed 93% head outgrowths, consistent with a repatterning of the nervous system occurring as DHs mature, (Figure 7B vii-ix). Taken together, these data reveal that the instructive influence of the CNS over the type of anatomical structures that will be formed after amputation changes over a much longer timescale than regeneration itself: in a mature double-headed animal, the CNS and tissue polarity remodel progressively (Figure 7C). Thus, the progressive remodeling of tissue polarity affects not only the epidermis but also the instructive aspects of the CNS.

## Discussion

Regeneration is a remarkable process because the activities of individual cells (migration, differentiation, proliferation, and shape change) need to be orchestrated toward a specific large-scale anatomical outcome. Order is simultaneously regulated at multiple scales and levels of organization, including tissues, organs, and whole-body axes. Planarian regeneration illustrates the ability of some systems to establish organs and physiological signaling systems in precise spatial coordination. This requires information about direction (alignment of planar polarity) as well as position (which organ to make at what location), but it is still very poorly understood how these distinct systems functionally interact. This is important not only for cell and developmental biology, but also for biomedicine. For example, planar polarity disruption is a known factor in carcinogenesis (Lee & Vasioukhin, 2008). Likewise, in the case of embryonic left-right patterning, significant birth defects occur when the cell-level alignment in organs such as the heart does not match the mirror-image inversion of body asymmetry (Delhaas, Decaluwe, Rubbens, Kerckhoffs, & Arts, 2004). Here, we exploited the planarian model system to reveal novel aspects of the linkage between axial and cell-level polarities, the role of the CNS in propagating the instructive signals that connect them, and the ways in which existing, well-formed tissues are remodeled in place to achieve the concordance between tissue polarity and organismal polarity.

We present evidence that planarians, which implement atypical body morphologies following interventions in normal regenerative signaling pathways, adjust the polarity of their existing tissues to align with these new morphologies on long timescales. DH planarians possess, in addition to their normal head positioned on the original anterior end, a secondary head positioned on the original posterior. This system gives us the opportunity to explore how planar cell polarity signaling, as read out by cilia orientation (Vu et al., 2018; Wallingford, 2010), changes in response to changes of overall organismal polarity. We show that cilia beat direction reverses in the portion of the double-headed animals that switches from a posterior to an anterior identity over a period of around 40 days. This long repatterning timeframe places this process outside of the normal regenerative timeframe commonly explored in studies with planarians, highlighting the importance of monitoring long-range remodeling events past the time when the damaged or missing tissues have been repaired or replaced.

The orientation of cilia is usually set during development or regeneration and is assumed to not to change once established. Here we have identified a system in which the dynamic reorientation of cilia induced by physiological signals, can be explored, thanks to the flexibility of the planaria body plan. The formation of new cells can be efficiently inhibited in planaria via irradiation, resulting in the death of the stem cells, while the non-dividing somatic tissue survives. We found that inhibition of new cell formation did not impact the reorientation of cilia, indicating that the reorientation of the cilia likely takes place in pre-existing multiciliated cells.

Exposure to the actin depolarizer Cytochalasin D speeds up the cilia reorientation process. We hypothesize that this effect may be due to a loosening of the actin cortex through depolymerization, allowing the basal bodies, whose attachment determines cilia beat orientation, to change their orientation easier, potentially by detaching from the membrane and reattach in the correct orientation. While the observed effect on cilia reorientation by cytochalasin D-driven actin depolymerization may be indirectly mediated via a number of different mechanisms, the crowded actin cortex in which the basal bodies are embedded offers a tantalizing explanation for this observation, where a reduction of cortex thickness or density may make reorientation of the cilia basal bodies easier. It is established, both in *Xenopus* embryos and developing mouse airway epithelia, that the actin cortex interacts with basal bodies to mediate basal body attachments and cilia orientation in development (Pan, You, Huang, & Brody, 2007; Panizzi, Jessen, Drummond, & Solnica-Krezel, 2007; Werner et al., 2011). Our data on actin depolymerization suggests that reorientation of planarian cilia during DH repatterning may follow a similar molecular process as the original establishment of cilia orientation in these developmental systems, giving subcellular level insights into the process driving reorientation of cilia.

We show that the reorientation of tissue is linked to the organism-scale polarity via signals from the brain which may be transmitted along the ventral nerve cords (VNCs), given that removal of the heads and lateral incisions leading to bisection of the VNCs affect the speed and extent of the repatterning similarly. We cannot rule out that other structures transmit the signaling factors driving repatterning, such as gap-junction-linked epidermal cells or the musculature which spans the length of the body, until further tools become available allowing us to identify signaling factor and its transport mechanism in more detail. The requirement for signals from the brain makes a transport of signaling factor along the VNCs a likely solution, as no transfer of the signal between different cell types would be necessary. The formation of an additional brain in the double-headed animal may provide an intrinsic signaling source to drive the required repatterning through the nervous system spanning the body. Our finding that bisection of the brain, without removal of tissue, results in the same magnitude of repatterning inhibition as removal of the entire head, suggests that the signal from the brain that drives repatterning relies on the intactness of the brain and is therefore likely to be a consequence of overall brain function or requires transport across the brain. Consistent with this idea, the overall size of the brain does not appear to influence in the rate of repatterning as long as it is intact, as indicated by irradiated animals which repattern normally even though they have very differently sized brains.

While signals from the brain are the major drivers of repatterning, a small amount of repatterning continues to happen after removal of both heads, which can be interpreted either as signal sent before decapitation continuing to be put into effect, or as a rudimentary repatterning system capable of driving limited repatterning via a different mechanism. This secondary repatterning mechanism can be speculated to consist of local PCP signaling, which is transmitted from cell to cell, leading to slower repatterning than that mediated by a neural path. We have hypothesized elsewhere that neural connections may have evolved in part to provide an optimized version of pre-neural signaling events operating in pattern regulation (Fields, Bischof, & Levin, 2019), so perhaps the repatterning that occurs in the absence of a head is an example of less efficient, ancestral non-neural patterning.

We find that during DH maturation, the midpoint is not fixed, given that the collision zone can be shifted across the midline through continuous removal of the primary head. While we were unable to keep the worms alive long enough to test whether full reversal of cilia beat direction across the entire animal is possible, our data suggest that it is. Our data illustrate that there is no secondary, underlying signal setting the midpoint of the epithelium but that this midpoint solely arises in normal DHs from the balance of signaling forces from the two heads.

At the same time, there appears to be a window of time after which the orientation of the cilia is set and can no longer be affected by removal of the animal’s head, as demonstrated by the lack of change in the position of the collision zone in mature DHs that had one head removed. Similarly, signals from the brain do not appear to be important for maintaining cilia orientation in single-headed animals. This suggests that there is a window of plasticity, during which signals from the brain can fully determine the polarity of the epithelium, but that once this window closes, cilia orientation is set, likely until changes induced by the regeneration of damaged or lost tissue occurs. This observation is reminiscent of data in developmental systems, were cilia orientation and tissue polarity are is set in certain developmental timeframes (Boisvieux-Ulrich, Sandoz, & Allart, 1991; Tung & Yeh-Tung, 1940; Twitty, 1928). What signals determine this plasticity in planarians remain to be addressed in future work.

Importantly, the link between global organ distribution and underlying tissue polarity exists in the nervous system too, not only in the epidermis. Our previous work showed how the global transport direction of the nervous system determines regenerative outcomes (Pietak et al., 2019). Here we observed that in maturing DHs, the symmetry point of the transport field of the nervous system shifts progressively to the middle of the animal, as revealed by the distinct regenerative outcomes of the same cuts along the body axis in immature and mature DHs. This suggests that the nervous system changes its overall polarity to adapt to the new, artificially induced morphologies. This is an extreme example of neuronal plasticity, opening new questions about what signals drive the changes in neural polarity and how correct and functional connections can re-form without interfering with CNS functions necessary for the animal to survive and continue its normal activity.

This work uses the dynamic body plan of the planarians as a way to explore multi-scale polarity *in vivo*. While organ structures are formed by the activity of cells, this bottom-up, self-assembly mechanism is complemented by top-down controls in which axial polarity drives changes of tissue-level directionality. The dynamic regeneration and repatterning of adult tissues in planarians offer an exciting addition to investigations of polarity in developmental systems. The mechanism of repatterning of the ciliated epithelium under the control of signals from the brain further highlights the importance of the nervous system as an overarching organizing system to coordinate and set polarity in regeneration and repatterning. At the same time the polarity control by the nervous system may be a bi-directional process, in which the nervous system acts both as a mediator of remodeling signals to the epithelium and at the same time is the target of remodeling. Beyond planarians, these findings suggest approaches that exploit brain-like signaling to understand and manipulate multi-scale order in bioengineering and developmental contexts.

## Materials & Methods

### Planaria colony care

Planaria of the species *Dugesia japonica* were maintained in a colony at 13 °C in Poland Spring water, fed once a week with calf liver paste and cleaned twice a week, as described in (Oviedo, Nicolas, Adams, & Levin, 2008). Animals were maintained at 13 °C for the time course of the entire experiment to prevent fissioning and were therefore taken from the cold-adapted colony at 13 °C, which is continuously kept at this temperature. Animals were starved for 1 week before use in experiments and for the course of the entire experiment to reduce variability due to metabolic state.

### Double-head induction

Double-headed planaria were generated by excision of pharynx and pre-tail fragments from planaria. These fragments were placed in 127 μM Octanol for 3 days before the solution was replaced with Poland Spring water and allowed to regenerate at 20 °C (Durant et al., 2017). Double-head regenerative outcomes were scored 7 days after cutting by the presence of a head, marked by at least one eye spot, on either end of the fragment. Double-head animal age was calculated based on the original cut, i.e. freshly regenerated DHs when they are selected, are here termed Day 7 DHs.

### Flow assay

Flow assays were performed based on original experiments in (Rustia, 1925). To detect cilia-driven flow, planaria were placed in a small dish with water, ventral-side up, so that they attached to the water surface. Carmine powder (Sigma-Aldrich, Darmstadt, Germany) was sprinkled on top of the worms. Powder and slime build-up was repeatedly removed using a paint brush. Movement of the powder particles was recorded on a Nikon AZ100M Multizoom Macroscope with a 0.5x objective and side-illumination from a Volpi IntraLED lightsource, using an Andor DL-604M camera. Movement was recorded at a frame rate of 100 ms. Animals were observed until powder accumulation reflecting cilia-driven flow had happened at least twice. Animals were washed 3 times in Poland Spring water to remove remaining carmine powder. Animals were maintained at 13 °C for duration of the repatterning assessment and were not fed.

Position of the collision zone was measured in ImageJ, where worm length and length from second head to midpoint of the powder collection point was measured. Both values were measured in worms which had even extension across the entire length of their body.

Particle image velocimetry (PIV) was performed using Matlab (Mathworks, Natick, MA) with the PIVlab program (Thielicke & Stamhuis, 2014, 2019). For the analysis, 2 sec segments were selected in which the worm exhibited little muscle movement and PIV was calculated in an interrogation window of 64 x 48 x 32 pixels for subsequent passes of the program, run on all frames, smoothed and all frames in the 2 sec segment were averaged to give flow pattern.

### Animal manipulations

Animals were irradiated using Nordion Gamma Cell 1000 Irradiator with a Cesium-137 radiation source with a dose of 200 Gy achieved via 30 min exposure at Day 7, unless otherwise specified.

Primary and secondary head identity was determined in animals at Day 7 based on development of head structures (eye size, pigmentation and auricle development) as well as overall movement pattern of the animal.

Decapitations were performed at the base of the head, directly underneath the auricles, at Day 7. Lateral cutting was performed at the respective positions between the two heads in animals placed ventral side up, so that ventral nerve cords were visible and cut as far across the body as necessary to sever both nerve cords, or to the midpoint for single VNC cutting. Nerve cord deviation was performed by cutting perpendicular to the head-head axis at the given point up to the midline of the worm and then continued along the midline, as described in (Pietak et al., 2019). In both cases cuts were reinforced every day for 7 days and worms were maintained at 13°C during cutting and regeneration. Regenerative outcomes were scored at Day 14.

Half-head removal, both lateral and perpendicular, was performed in DHs at Day 7 following irradiation. The secondary head of the animals was cut in all these cases. For lateral head removal a cut was placed downward between the eyes and then perpendicular at the base of the head to remove half of the head. For the perpendicular half-head removal, the cutting plane was perpendicular to the body axis and set at the plane of the eyes. Cuts were confirmed using synapsin staining.

Cuts bisecting the brain were performed in DHs at Day 7 following irradiation, targeting the secondary head. The cut was placed downwards between the eyes to the base of the head. The cut was re-enforced for 3 days following the initial cut.

Internal tissue removal was performed using square glass capillaries of a 1 mm^2^ inner diameter and 0.1 mm wall thickness (VitroCom, Mountain Lakes, NJ), using a fine paint brush to remove the cut tissue from the middle of the animal.

Double-headed animals at Day 7 and Day 49 were cut perpendicular to the head-head axis either at the midpoint between the two heads, halfway between the midpoint and the head, i.e. at 25% body length, and directly at the base of either the primary or secondary head. Fragments regenerated at 20°C for 14 days before scoring regenerative outcomes by counting number of single-headed and double-headed regenerates.

Brightfield pictures of animals were taken on a Nikon SMZ1500.

### Drug treatments

Chemicals used to disrupt the cilia repatterning process were applied to freshly regenerated DHs at Day 7 and maintained throughout the repatterning process with weekly replacement of drugs, unless otherwise noted. Inhibitors used were Cytochalasin D (Sigma) at 1 μM final concentration from stock dissolved in Ethanol, Serotonin (Sigma) at a final concentration of 0.5 mM from a stock dissolved in H_2_O, Fluoxetine (Sigma) at a final concentration of 2 μM from a stock dissolved in DMSO, and 5,7-Dihydroxytryptamine (ThermoFisher Scientific, Waltham, MA) at a final concentration of 75 μM from a stock dissolved in H_2_O.

200 proof Ethanol (VWR, Radnor, PA) was used at 3% in Poland spring water to briefly disrupt cilia (Stevenson & Beane, 2010) by incubating for 1 h before washing out.

### Fixation and Immunohistochemistry

Animals were fixed at the described time point using an 2% HCl treatment for 2 min followed by incubation in Carnoy’s fixative (60% Ethanol, 30% Chloroform, 10% Acetic Acid) on ice for 2 hours, before washing with Methanol at −20 °C and overnight bleaching with 10% H_2_O_2_ in Methanol. Fixed samples were stained using the VSI InSitu Pro robot (Intavis, Cologne, Germany), specifically rehydrated in PBSTx (1x PBS with 0.3% Triton X-100), blocked in 10% Goat serum in PBSTx+B (1x PBS with 0.3% Triton X-100 and 10% BSA) for 6 hours, stained in primary and secondary antibodies (10 hours at 4 °C for both) and washed with PBSTx (2x 20 min and 1x 1 hour). The primary antibody used were - for synapsin, mouse-anti-synapsin (SYNORF1) antibody at 1:50 (Developmental Studies Hybridoma Bank (DSHB), University of Iowa), for dividing cells, rabbit anti-phospho-Histone H3 (Ser10) clone MC463 (Millipore Sigma) at 1:250, and for cilia, mouse-anti-acetylated tubulin (Sigma) at 1:1000. Secondary antibody used was goat-anti-mouse IgG (H+L) Cross-Adsorbed-Alexa555 (ThermoFisher) at 1:400 and goat-anti-rabbit-HRT 1:100, with TSA-Alexa488 amplification for the phosphor-Histone H3 antibody. All antibodies were diluted in PBSTx+B. Anti-SYNORF1 was deposited to the DSHB by E. Buchner (DSHB Hybridoma Product 3C11 (anti-SYNORF1)) (Klagges et al., 1996).

Samples were mounted in Vectashield^®^ hard set mounting medium (Vector Laboratories, Burlingame, CA) and imaged on a Leica SP8 confocal (Leica, Mannheim, Germany) with HyD detector, 488 nm and 552 nm diode light source, and 10x NA=0.4 (Leica HC PL APO CS2) or 25x water-immersion NA=0.95 (Leica HC Fluotar L) objectives.

### Image analysis, data analysis and statistical information

Sample sizes were chosen in accordance with standards of the field. Repeat numbers and sample sizes are reported in the respective figure legends. Animals taken from the same colony unit and treated at the same time and investigated in parallel are considered technical replicates, while animals from a different colony unit investigated at a different time represent biological replicates. Animals were randomly selected out of large colony units and randomly assigned into treatment groups. Data analysis was performed blinded.

All image analysis, measurements and image post-processing were performed in FIJI ImageJ (Rueden et al., 2017; Schindelin et al., 2012). Data analysis and plotting was done in GraphPad Prism V8.0 (GraphPad, San Diego, CA). All datapoints are plotted where reasonable, otherwise center and dispersion measures are defined in the respective figure legends. All data was included in all analyses. All plotted data can be found in Supplementary Dataset 1.

Unless otherwise specified, multiple t-tests with false discovery rate (FDR) approach using the two-stage step-up method of Benjamini, Krieger and Yekutieli with an FDR = 1 %, not assuming equal SD, was performed to analyze all datasets. All other datasets were analyzed using a Welsh’s two-tailed t-test. Significance threshold was set to p<0.01. Statistical test data are given in respective Supplementary Tables.

## Acknowledgements

The authors would like to thank Anna Kane, Joshua Finkelstein, Alexis Pietak, and all members of the Levin lab for thoughtful discussions on this project. We thank Nicolas Spitzer, Mansi Shrivastava, and James Monaghan for critical reading and comments on the manuscript. We thank Junji Morokuma and Hans Gonzembach for planaria colony maintenance. This research was supported by the Allen Discovery Center program through The Paul G. Allen Frontiers Group (12171), and by the National Institutes of Health Research Infrastructure grant NIH S10 OD021624.

## Author contribution

Conceptualization: Johanna Bischof and Michael Levin

Methodology: Johanna Bischof

Validation: Johanna Bischof and Margot E. Day

Formal Analysis: Johanna Bischof

Investigation: Johanna Bischof, Margot E. Day, Kelsie A. Miller, Joshua LaPalme

Resources: Michael Levin

Data Curation: Johanna Bischof, Margot E. Day

Writing – Original Draft Preparation: Johanna Bischof

Writing – Review and Editing: Johanna Bischof, Michael Levin, Margot E. Day, Kelsie A. Miller, Joshua LaPalme

Visualization: Johanna Bischof

Supervision: Michael Levin

Project Administration: Johanna Bischof

Funding Acquisition: Michael Levin

## Videos

**Video 1: Movement of a DH planarian at Day 7.** The two heads are not evenly regenerated, and the more fully regenerated head dominates the movement of the whole animal. Sped up 2x.

**Video 2: Movement of a DH planarian at Day 49.** The two heads are evenly regenerated, and movement is equally controlled by both halves of the animal, leading to frequent stalling. Sped up 2x.

**Video 3: Carmine powder flow assay in a single-headed animal.** Single-headed planarian showing unidirectional flow of the carmine powder across the whole animal. Sped up 20x, 171 frames at 100 ms frame rate.

**Video 4: Position of the collision zone changes over time in DH animals.** Flow in DH animals at Day 7, Day 14, Day 21, Day 28, Day 35 and Day 42, showing progressive movement of the collision zone across the posterior half of the animal. Sped up 5x, 26 frames at 100 ms frame rate.

**Video 5: Cilia driven flow in animals with internal tissue removal.** Flow in an DH animal following internal tissue removal at Day 7, flow imaged at Day 28. Asterisk marks site of healed wound. Sped up 5x, 26 frames at 100 ms frame rate.

**Video 6: Serotonin treatment blocks flow.** A) Untreated DH and B) DH treated with 500 μM Serotonin at Day 14. Sped up 10x, 101 frames at 100 ms frame rate.

**Video 7: Removal of heads impacts repatterning speed.** A) Control DH, B) DH with primary head removed showing further advanced collision zone compared to control, C) DH with secondary head removed showing less advanced collision zone compared to control, D) DH with both heads removed showing equally less advanced collision zone compared to control at Day 35. Sped up 5x, 26 frames at 100 ms frame rate.

**Video 8: Continuous removal of primary head allows repatterning across the midpoint.** DH with its primary head continuously removed with the collision zone positioned across the midline, as visible when the animal is fully extended, at Day 56. Sped up 5x, 51 frames at 100 ms frame rate.

**Video 9: Removal of head in a mature DH does not reposition the collision zone.** A) DH animal at Day 49 before treatment. B) DH animal irradiated and with one head removed at Day 49, imaged at Day 63. Sped up 5x, 51 frames at 100 ms frame rate.

**Video 10: Removal of head in SH animal does not impact ciliary orientation.** A) SH animal before treatment showing unidirectional flow. B) Ciliary driven flow in the same animal following irradiation, removal of head and incubation for 21 days. Sped up 10x, 82 frames at 100 ms frame rate.

**Video 11: Cilia-driven flow in an animal with a lateral cut.** Animal cut laterally following irradiation at Day 7 in the secondary half of the body, showing collision zone positioned behind the cut. Sped up 5x, 51 frames at 100 ms frame rate.

## Supplemental Figures

**Figure S1:**
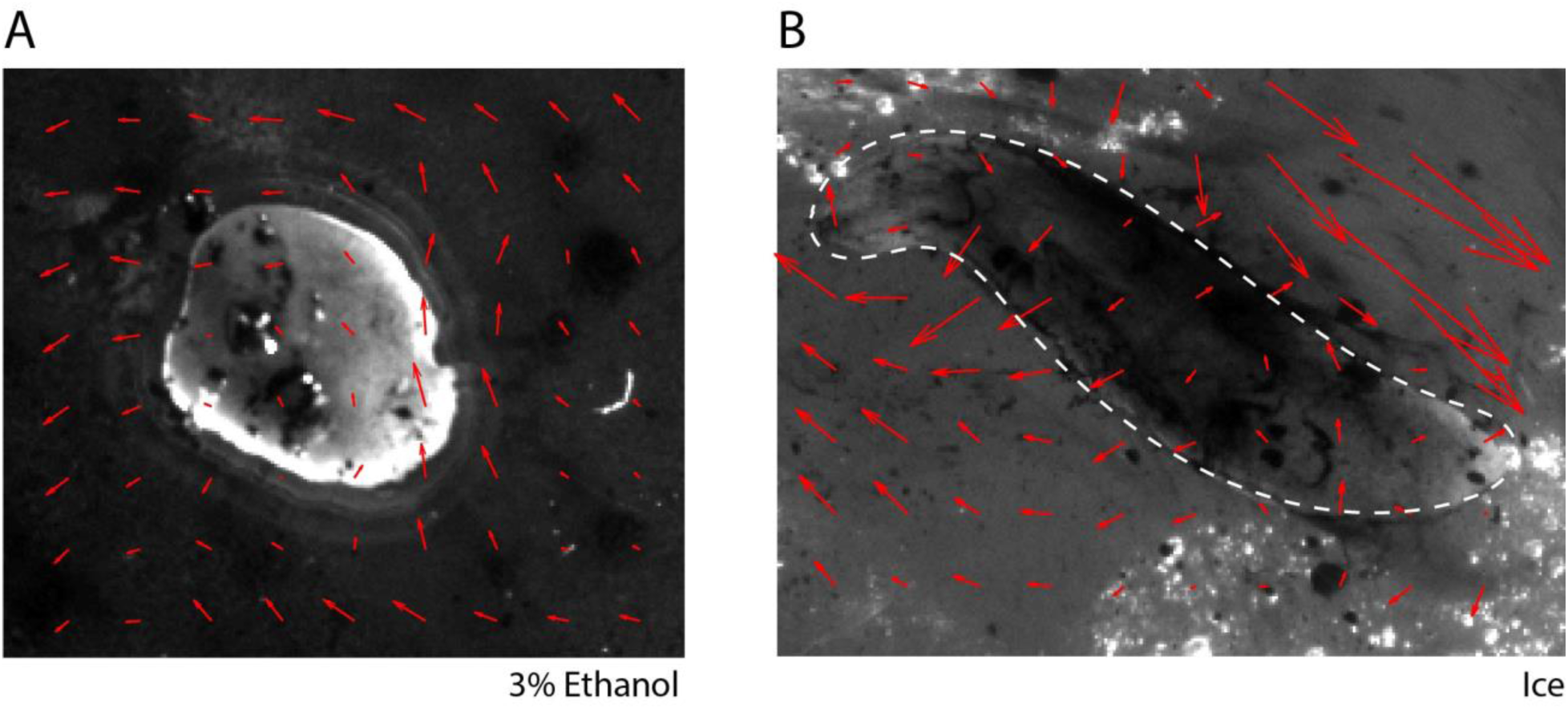
Controls for carmine-powder flow assay. A) Animal treated with 3% Ethanol for 1 hour before testing shows no flow, as visualized by tracking of particle movement using particle image velocimetry (PIV) (red arrows). B) Animal maintained on ice to reduce temperature shows no flow along the animal axis (outlines by dashed line), as visualized by tracking of particle movement by PIV (red arrows).

**Figure S2:**
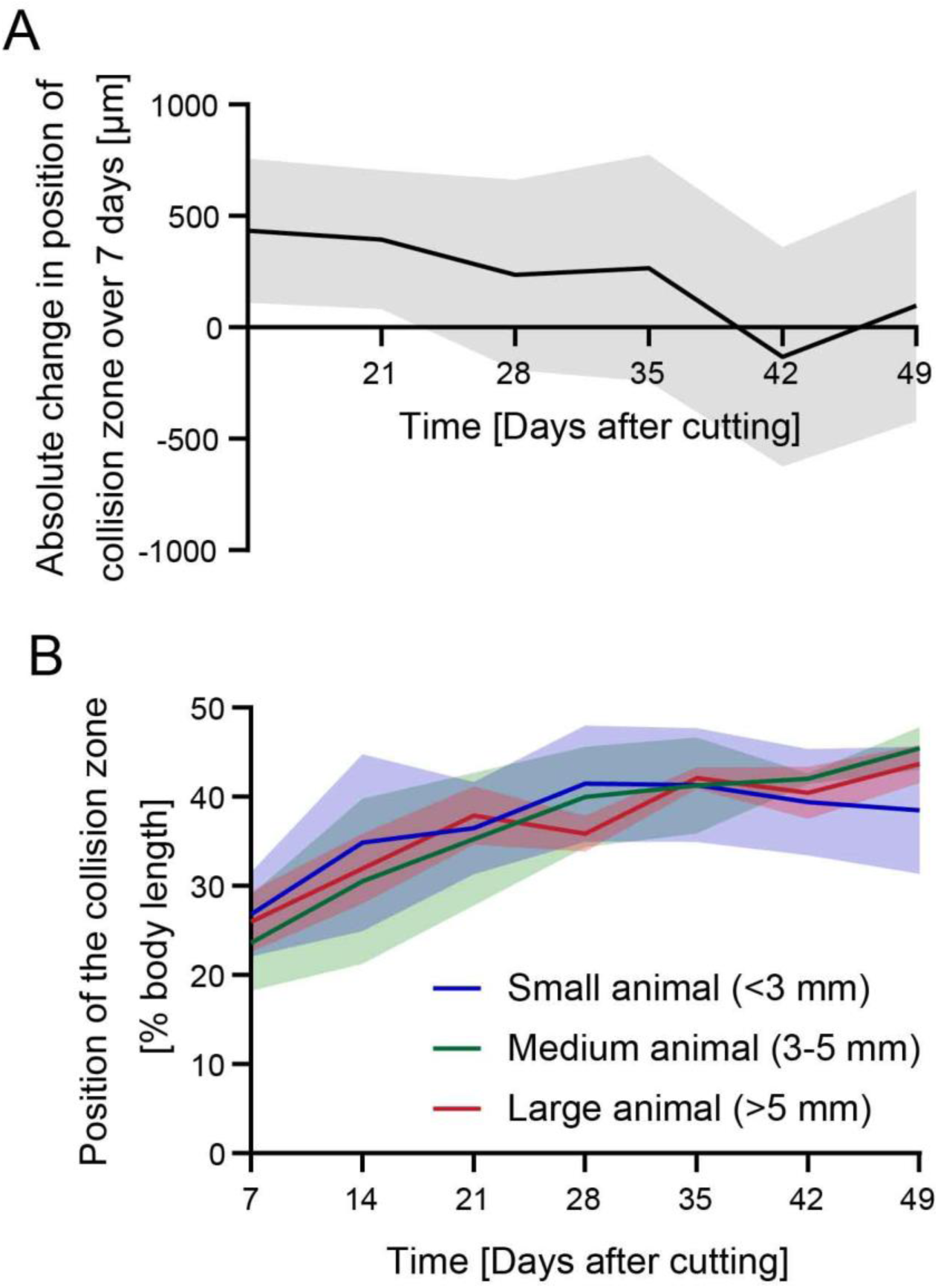
Progression of cilia reorientation. A) The absolute change in the position of the collision zone is relatively even from Day 7 until Day 35, when it slows down and stalls following D42. N=24. B) Reorientation of the cilia position occurs at the same relative speed in animals of different sizes, suggesting scaling of the repatterning process. N=4. Plotted is mean of all samples, with standard deviation represented as shaded area.

**Figure S3:**
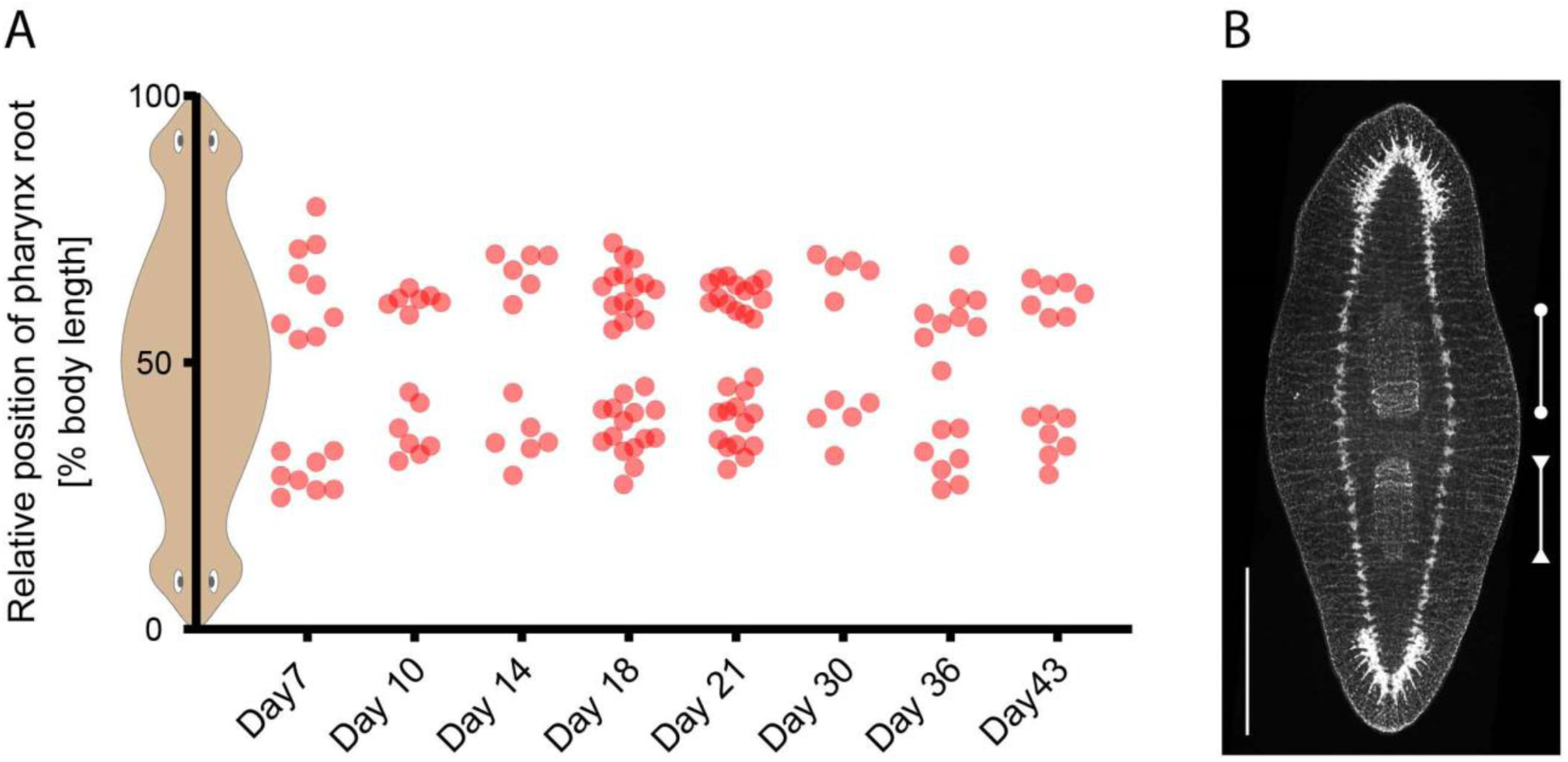
Position of the pharynges in double-headed animals at different ages. A) Position of root of the primary and secondary pharynx in DH animals at different timepoints from Day 7 to Day 43, plotted as position along the body length of the double-headed animal. B) Synapsin antibody stain in a double-headed animal at Day 14, showing the pharynx in the primary half of the animal (bar with circular ends) and the pharynx in the secondary half of the animal (bar with triangular ends), illustrating position of the two structures. Scale bar 1mm.

**Figure S4:**
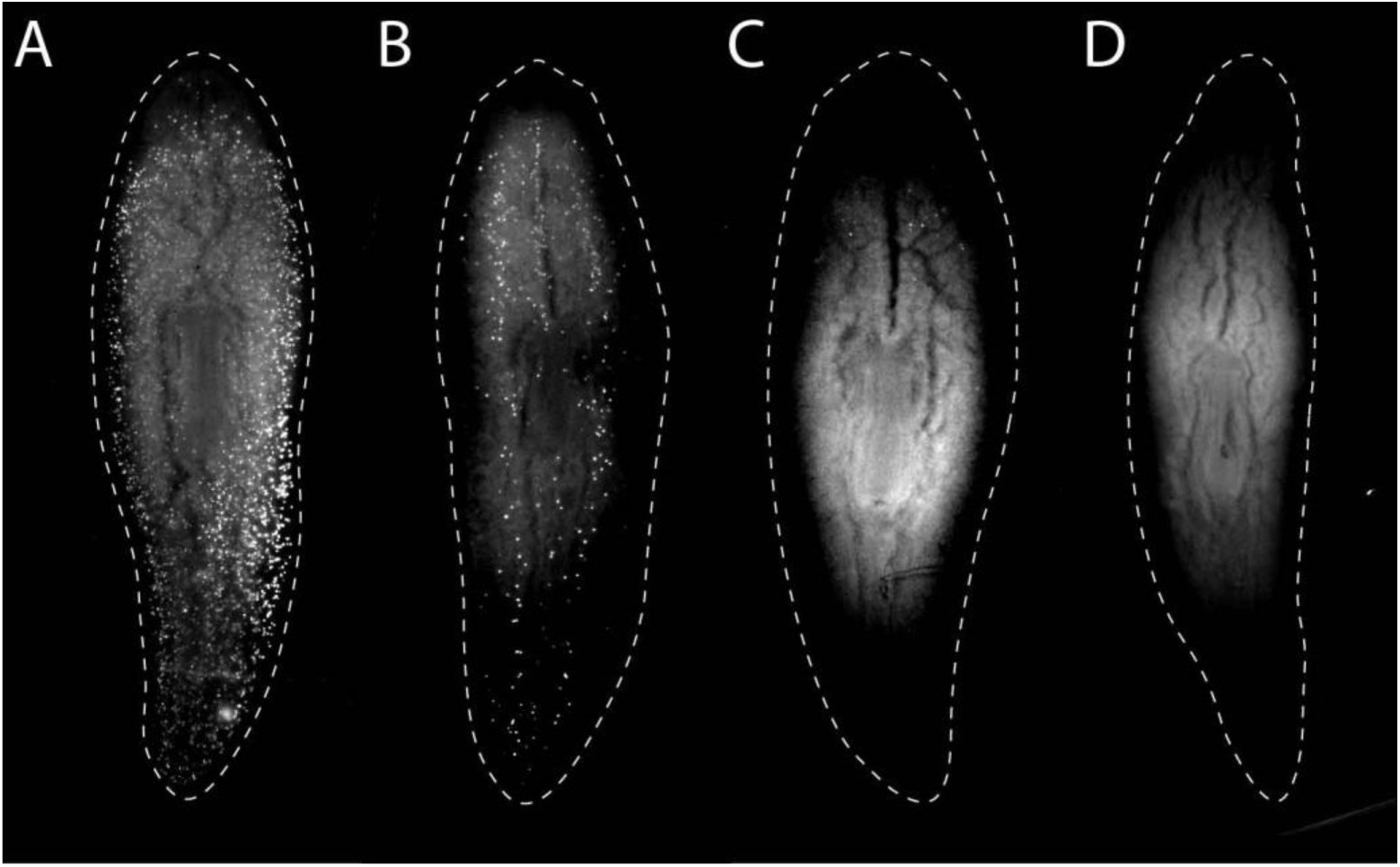
Depletion of dividing cells following irradiation. A) Unirradiated control animal, and animals B) 1 day, C) 3 days and D) 7 days after irradiation with 200 Gy. Stained with anti-phospho-Histone H3 (Ser10).

**Figure S5:**
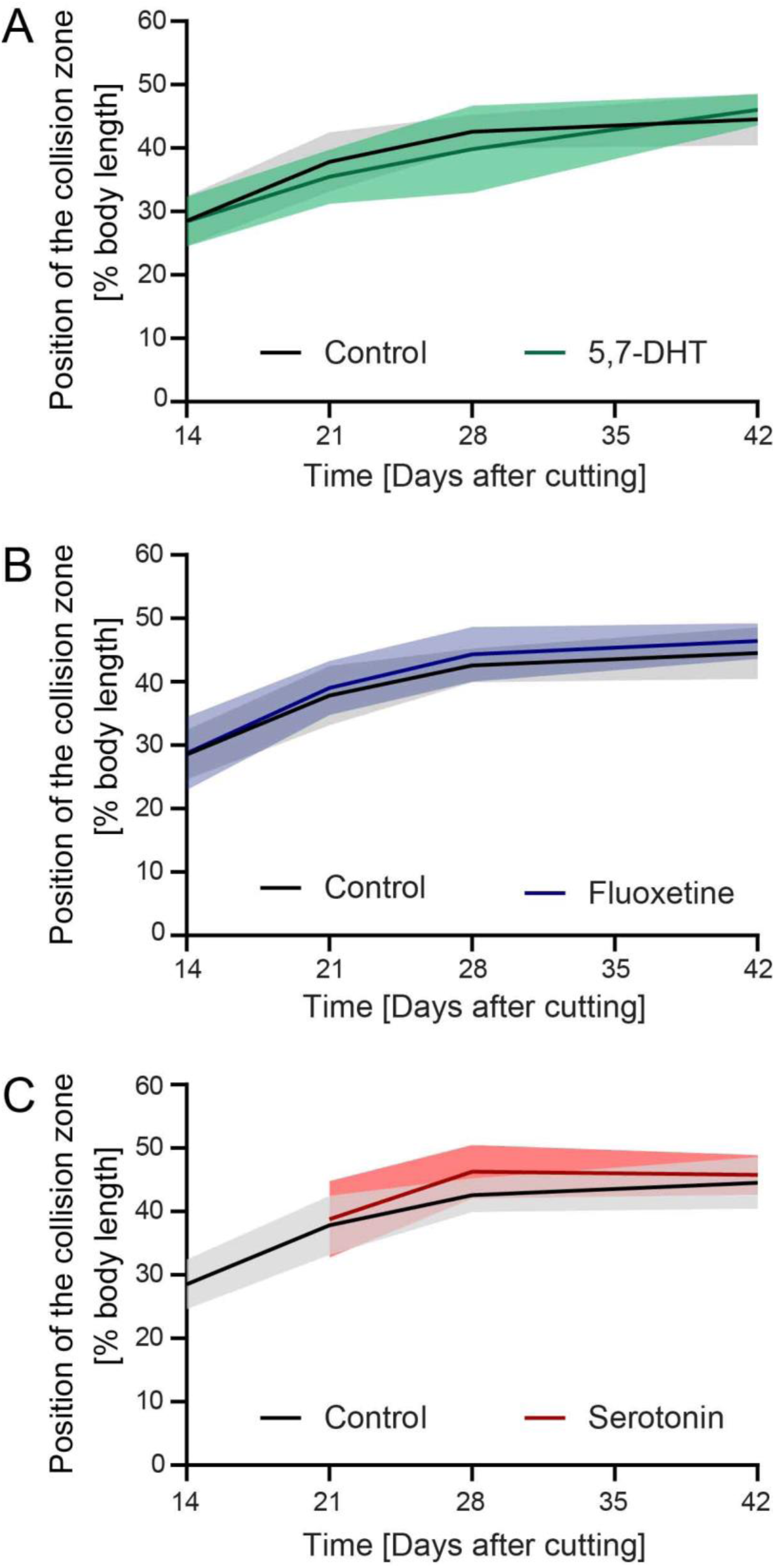
Treatments disrupting serotonergic neurons do not affect cilia reorientation. A) DHs treated continuously from Day 7 onwards with 75 μM 5,7-Dihydroxytryptamine (5,7-DHT) (green), a serotonergic neurotoxin, showed no difference in repatterning speed compared to controls (black). B) DHs treated continuously from Day 7 onwards with 2 μM Fluoxetine (blue), a serotonin reuptake inhibitor, showed no difference in repatterning speed compared to controls (black). C) DHs treated from Day 7 till Day 14 with 500 μM serotonin (red) showed no difference in repatterning speed compared to controls (black) after washout of the drug. No flow was detectable during serotonin incubation. N=12. Plotted is mean of all samples, with standard deviation as shaded area, all experiments with their age-matched controls.

**Figure S6:**
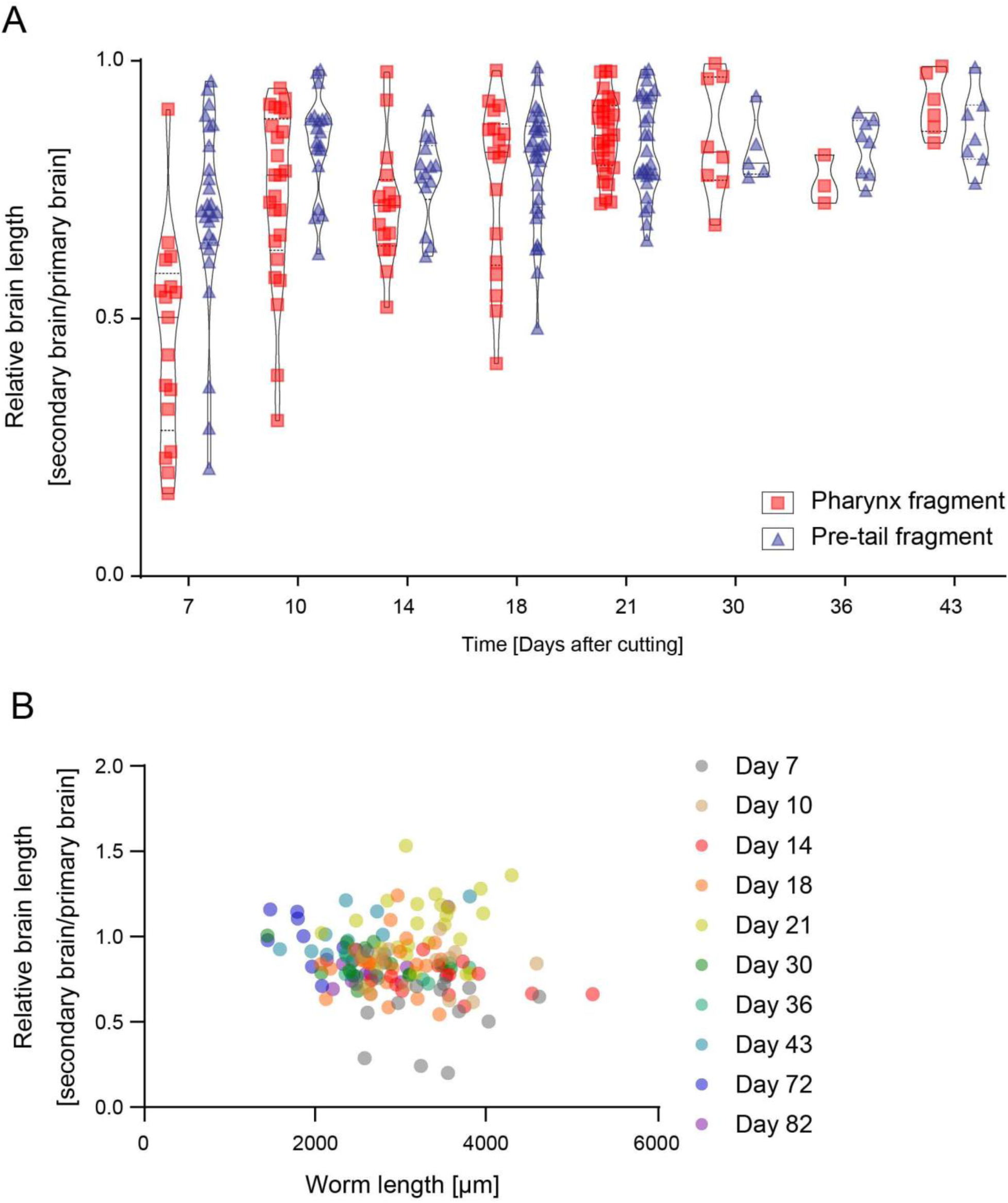
Brain size differences in DHs do not correlate with fragment origin or size of animal. A) Relative size of the brain in the primary and secondary head in DHs generated either from pharynx (red square) or pre-tail fragments (blue triangle) from Day 7 to Day 43, showing no pronounced difference in the distribution of the data at each timepoint. B) Relative brain length of the primary and secondary brain over time plotted against worm size, showing no correlation with worm length.

**Figure S7:**
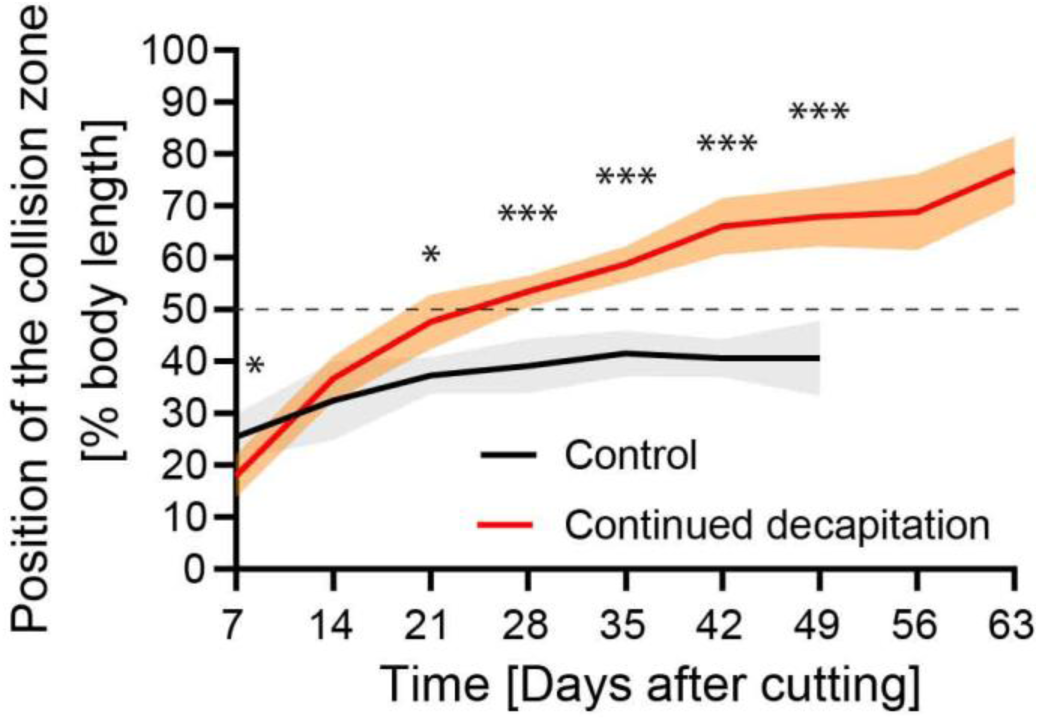
Serial removal of the primary head allows repatterning across the midline. Repatterning in animals which had their primary head removed every second day after Day 7 (red), compared to controls (black), showing significant difference in repatterning speed and advancement of collision zone far above the 50% boundary. N=12. Plotted is mean of all samples, with standard deviation as shaded area, all experiments with their age-matched controls.

**Figure S8:**
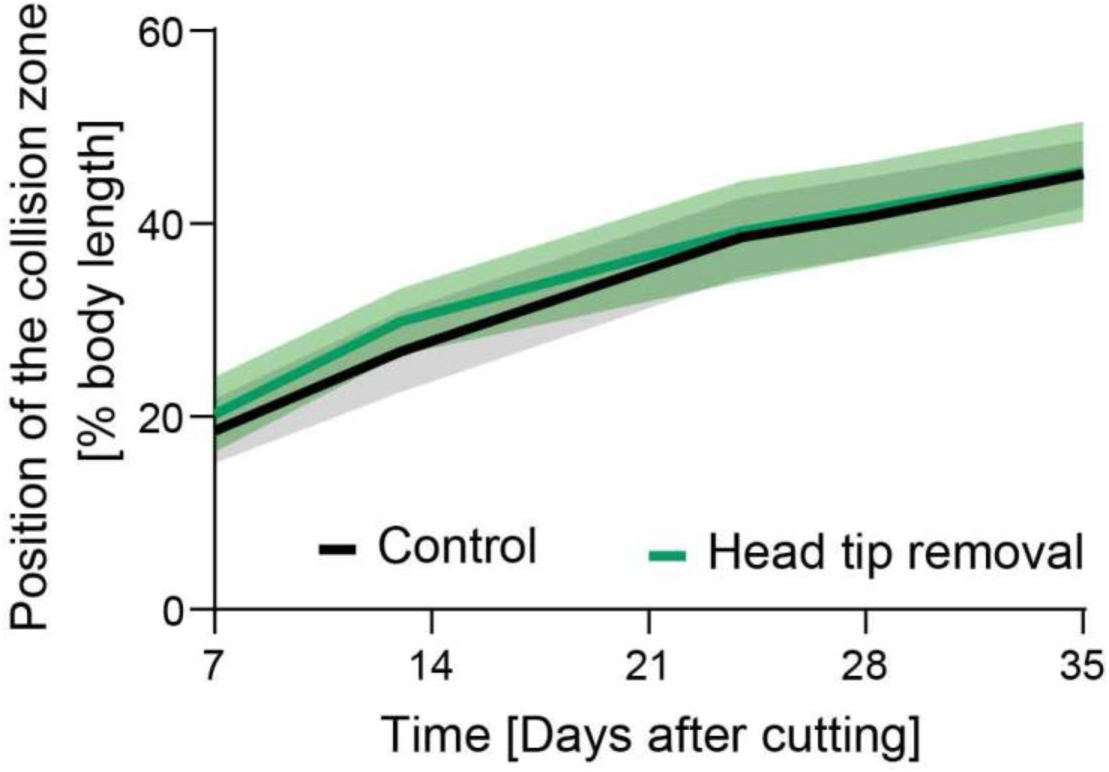
Removal of the head tip does not impact repatterning. DHs irradiated and with the head tip of the secondary head removed at Day 7 showed no difference in repatterning speed compared to controls. N=12. Plotted is mean of all samples, with standard deviation as shaded area, all experiments with their age-matched controls.

**Figure S9:**
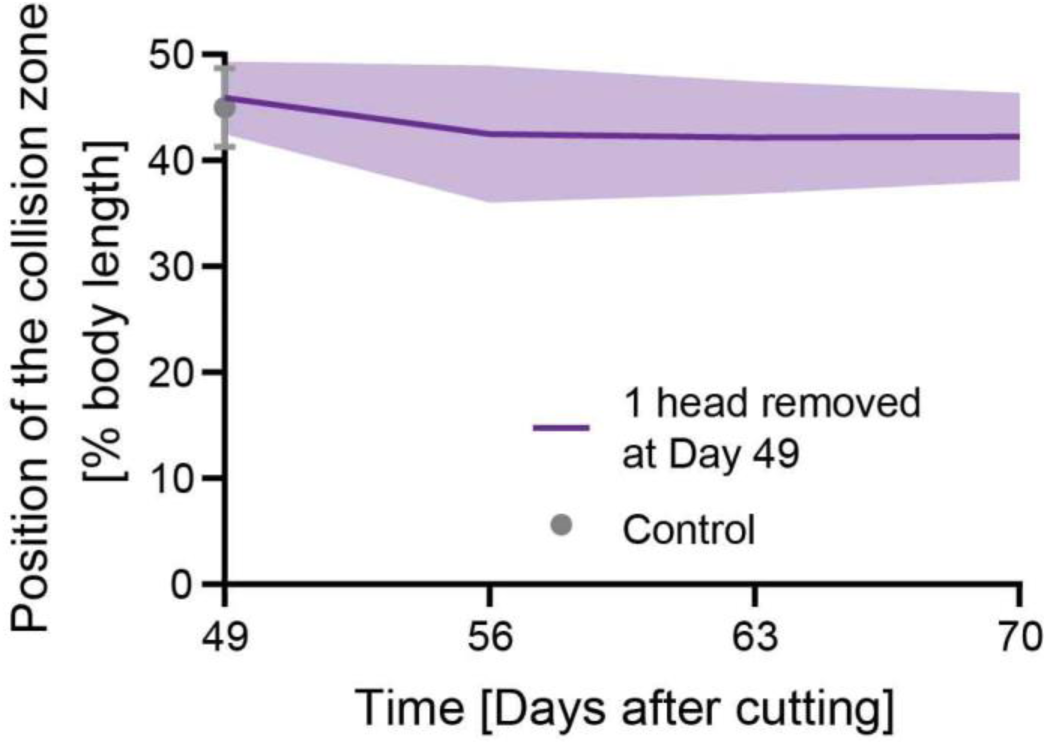
Removal of one head in mature DHs does not impact position of collision zone. Position of the collision zone does not change in DHs, which had one of their heads removed at Day 49 following irradiation, tracked over 21 days following head removal. N=12. Plotted is mean of all samples, with standard deviation as shaded area. Grey point with error bars represents control sample at Day 49.

**Figure S10:**
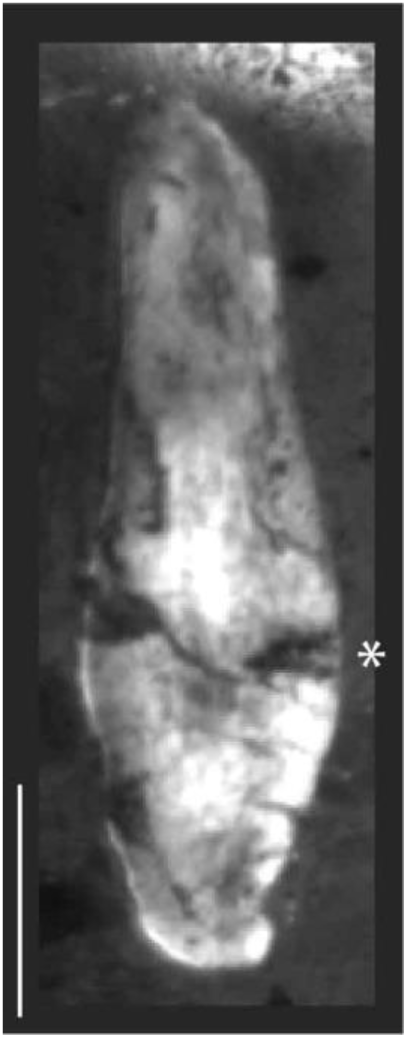
DHs with one side of the secondary head removed show no significant angling of the collision zone. DHs were irradiated and half of the head was removed laterally at Day 7 and flow was tracked. Image of worm at Day 28.

**Figure S11:**
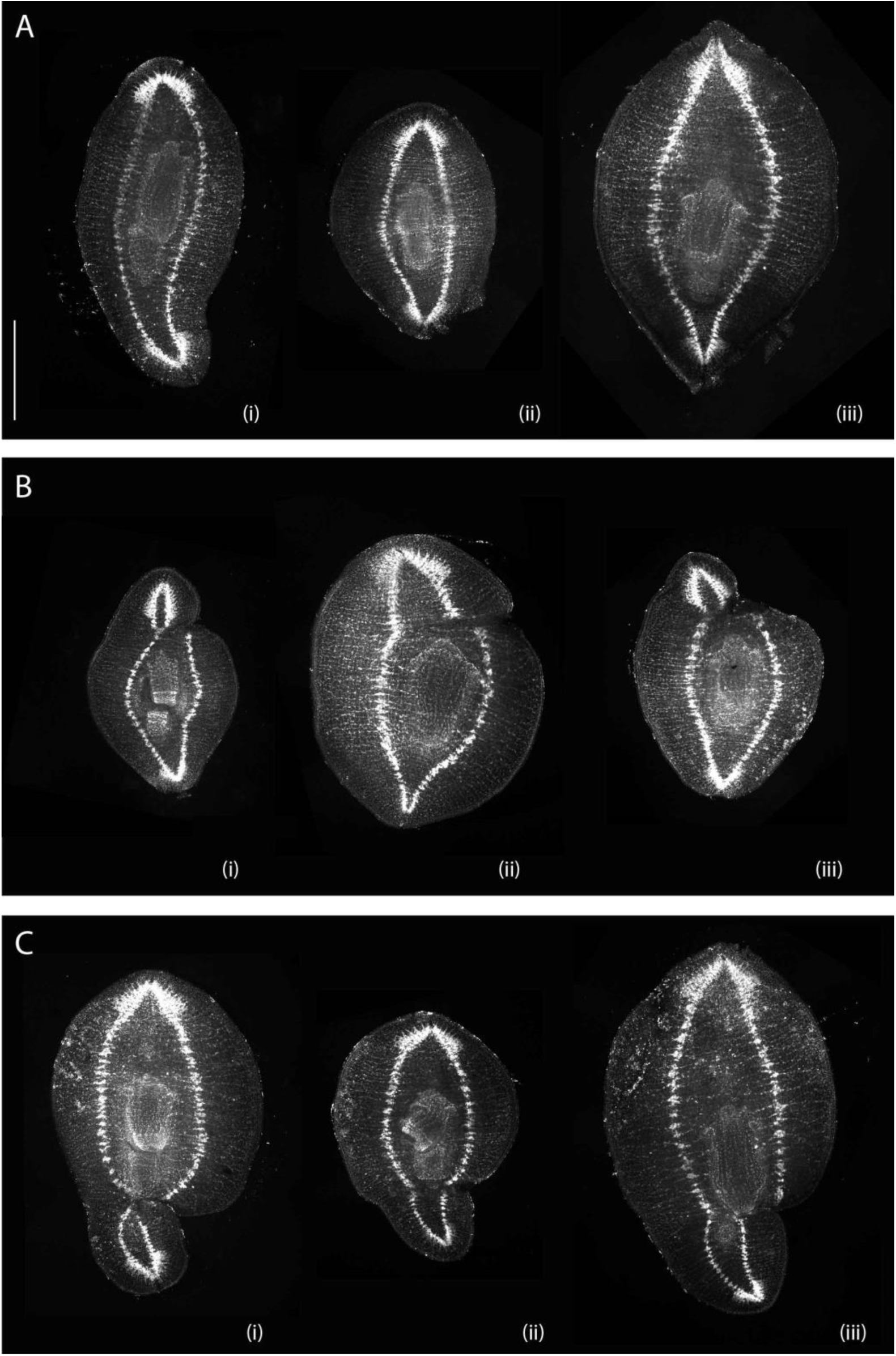
Further examples of laterally cut animals stained with synapsin antibody. A) Control animals. B) DHs with cut underneath the primary head, showing diversity of cut and healing outcomes. C) DHs with cut underneath the secondary head. All animals were irradiated and cut at Day 7 and fixed at Day 28. Scale bar 1 mm.

## Supplemental Tables

**Supplementary Table 1:** Internal tissue removal, statistical analysis data.

| Time point | P value | Mean of Control | Mean of Large Wormholes | Difference | SE of difference | t ratio | df | q value |
| --- | --- | --- | --- | --- | --- | --- | --- | --- |
| Day 21 | 0.943328 | 34.78 | 34.99 | -0.2128 | 2.953 | 0.07209 | 18.00 | 0.952761 |
| Day 32 | 0.406620 | 41.80 | 43.35 | -1.556 | 1.831 | 0.8498 | 18.00 | 0.616029 |
| Day 39 | 0.230430 | 46.80 | 45.47 | 1.324 | 1.064 | 1.244 | 17.00 | 0.616029 |

**Supplementary Table 2:**
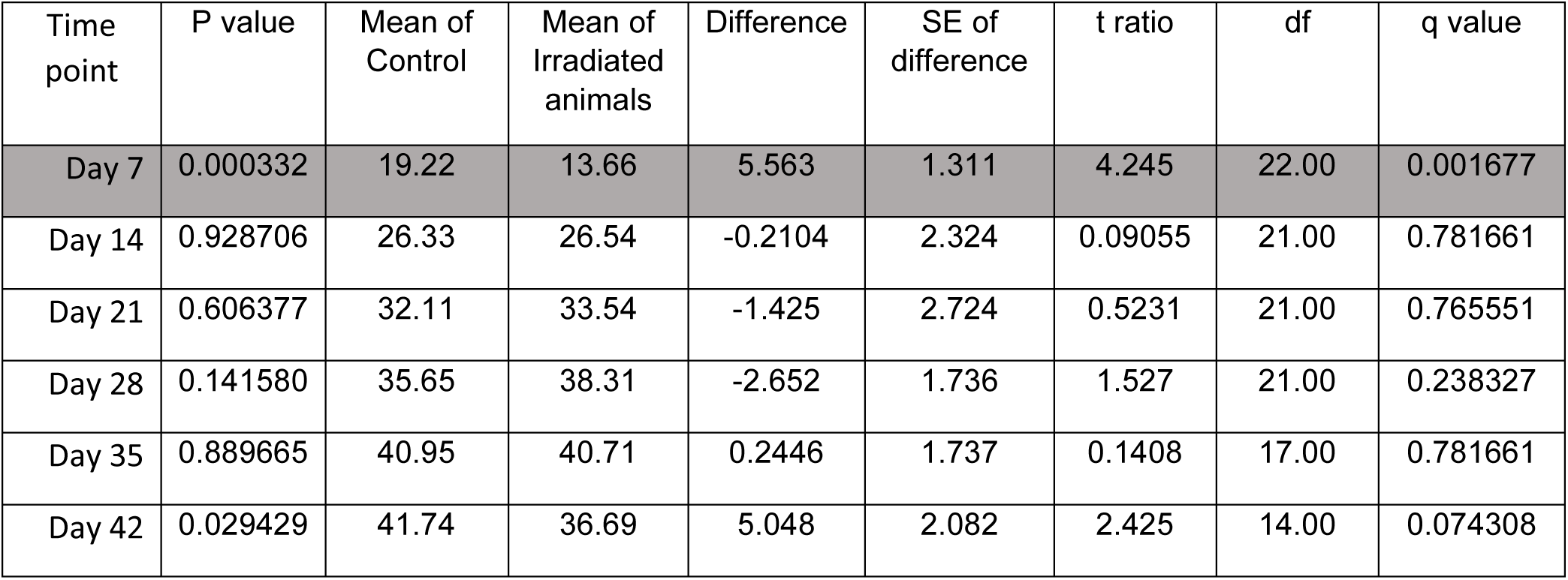
Irradiation, statistical analysis data.

| Time point | P value | Mean of Control | Mean of Irradiated animals | Difference | SE of difference | t ratio | df | q value |
| --- | --- | --- | --- | --- | --- | --- | --- | --- |
| Day 7 | 0.000332 | 19.22 | 13.66 | 5.563 | 1.311 | 4.245 | 22.00 | 0.001677 |
| Day 14 | 0.928706 | 26.33 | 26.54 | -0.2104 | 2.324 | 0.09055 | 21.00 | 0.781661 |
| Day 21 | 0.606377 | 32.11 | 33.54 | -1.425 | 2.724 | 0.5231 | 21.00 | 0.765551 |
| Day 28 | 0.141580 | 35.65 | 38.31 | -2.652 | 1.736 | 1.527 | 21.00 | 0.238327 |
| Day 35 | 0.889665 | 40.95 | 40.71 | 0.2446 | 1.737 | 0.1408 | 17.00 | 0.781661 |
| Day 42 | 0.029429 | 41.74 | 36.69 | 5.048 | 2.082 | 2.425 | 14.00 | 0.074308 |

**Supplementary Table 3:** Short term ethanol treatment, statistical analysis data.

|  | P value | Mean of Control | Mean of 1h Ethanol | Difference | SE of difference | t ratio | df | q value |
| --- | --- | --- | --- | --- | --- | --- | --- | --- |
| Day 7 | 0.002248 | 25.42 | 18.37 | 7.057 | 2.029 | 3.478 | 21.00 | 0.015891 |
| Day 14 | 0.293892 | 32.44 | 29.32 | 3.121 | 2.899 | 1.077 | 21.00 | 0.478683 |
| Day 21 | 0.362489 | 37.33 | 34.53 | 2.801 | 2.975 | 0.9413 | 14.00 | 0.478683 |
| Day 28 | 0.088478 | 39.11 | 42.58 | -3.468 | 1.941 | 1.786 | 21.00 | 0.245426 |
| Day 35 | 0.989495 | 41.54 | 41.57 | -0.02368 | 1.776 | 0.01333 | 20.00 | 0.999390 |
| Day 42 | 0.104141 | 40.63 | 43.15 | -2.526 | 1.484 | 1.703 | 20.00 | 0.245426 |

**Supplementary Table 4:** 5,7-Dihydroxytroptamine treatment, statistical analysis data.

|  | P value | Mean of Control | Mean of DHT animals | Difference | SE of difference | t ratio | df | q value |
| --- | --- | --- | --- | --- | --- | --- | --- | --- |
| Day 14 | 0.947371 | 28.54 | 28.43 | 0.1089 | 1.630 | 0.06680 | 21.00 | 0.956845 |
| Day 21 | 0.209620 | 37.82 | 35.46 | 2.361 | 1.827 | 1.292 | 22.00 | 0.390030 |
| Day 28 | 0.209798 | 42.57 | 39.83 | 2.737 | 2.119 | 1.292 | 22.00 | 0.390030 |
| Day 42 | 0.289626 | 44.54 | 46.02 | -1.481 | 1.365 | 1.085 | 22.00 | 0.390030 |

**Supplementary Table 5:**
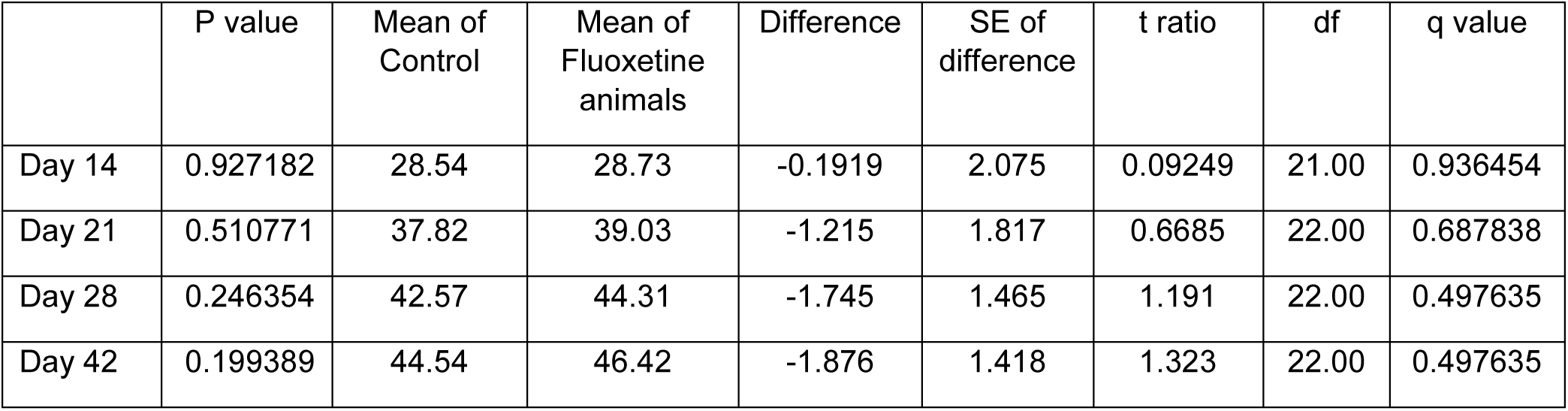
Fluoxetine treatment, statistical analysis data.

|  | P value | Mean of Control | Mean of Fluoxetine animals | Difference | SE of difference | t ratio | df | q value |
| --- | --- | --- | --- | --- | --- | --- | --- | --- |
| Day 14 | 0.927182 | 28.54 | 28.73 | -0.1919 | 2.075 | 0.09249 | 21.00 | 0.936454 |
| Day 21 | 0.510771 | 37.82 | 39.03 | -1.215 | 1.817 | 0.6685 | 22.00 | 0.687838 |
| Day 28 | 0.246354 | 42.57 | 44.31 | -1.745 | 1.465 | 1.191 | 22.00 | 0.497635 |
| Day 42 | 0.199389 | 44.54 | 46.42 | -1.876 | 1.418 | 1.323 | 22.00 | 0.497635 |

**Supplementary Table 6:** Serotonin treatment, statistical analysis data.

|  | P value | Mean of Control | Mean of Serotonin animals | Difference | SE of difference | t ratio | df | q value |
| --- | --- | --- | --- | --- | --- | --- | --- | --- |
| Day 21 | 0.685684 | 37.82 | 38.80 | -0.9823 | 2.388 | 0.4113 | 18.00 | 0.692541 |
| Day 28 | 0.026119 | 42.57 | 46.27 | -3.709 | 1.530 | 2.424 | 18.00 | 0.052761 |
| Day 42 | 0.484950 | 44.54 | 45.75 | -1.211 | 1.698 | 0.7131 | 18.00 | 0.653066 |

**Supplementary Table 7:** Cytochalasin D treatment, statistical analysis data.

|  | P value | Mean of Control | Mean of CytoD animals | Difference | SE of difference | t ratio | df | q value |
| --- | --- | --- | --- | --- | --- | --- | --- | --- |
| Day 7 | 0.167679 | 18.87 | 16.72 | 2.152 | 1.509 | 1.427 | 22.00 | 0.225808 |
| Day 14 | 0.899292 | 25.88 | 25.65 | 0.2237 | 1.746 | 0.1281 | 21.00 | 0.605523 |
| Day 21 | 0.000235 | 33.06 | 37.19 | -4.138 | 1.068 | 3.873 | 72.00 | 0.000474 |
| Day 28 | 0.000004 | 37.76 | 42.24 | -4.478 | 0.9002 | 4.974 | 78.00 | 0.000015 |
| Day 35 | 0.880184 | 40.81 | 40.52 | 0.2898 | 1.898 | 0.1527 | 20.00 | 0.605523 |
| Day 42 | 0.882505 | 41.65 | 41.85 | -0.1958 | 1.308 | 0.1497 | 20.00 | 0.605523 |

**Supplementary Table 8:** Primary head removal, statistical analysis data.

|  | P value | Mean of Control | Mean of Primary decapitation animals | Difference | SE of difference | t ratio | df | q value |
| --- | --- | --- | --- | --- | --- | --- | --- | --- |
| Day 7 | 0.677534 | 19.22 | 19.99 | -0.7705 | 1.828 | 0.4214 | 22.00 | 0.114052 |
| Day 14 | 0.007399 | 26.33 | 33.10 | -6.770 | 2.284 | 2.964 | 21.00 | 0.001495 |
| Day 21 | 0.000565 | 32.11 | 39.33 | -7.222 | 1.780 | 4.058 | 21.00 | 0.000285 |
| Day 28 | 0.001164 | 35.65 | 44.55 | -8.897 | 2.351 | 3.784 | 20.00 | 0.000392 |
| Day 35 | 0.000030 | 40.95 | 50.15 | -9.198 | 1.665 | 5.526 | 18.00 | 0.000031 |
| Day 42 | 0.003744 | 41.74 | 51.19 | -9.451 | 2.816 | 3.357 | 17.00 | 0.000945 |

**Supplementary Table 9:** Secondary head removal, statistical analysis data.

|  | P value | Mean of Control | Mean of Secondary decapitation animals | Difference | SE of difference | t ratio | df | q value |
| --- | --- | --- | --- | --- | --- | --- | --- | --- |
| Day 7 | 0.662416 | 19.22 | 18.61 | 0.6148 | 1.389 | 0.4425 | 22.00 | 0.223013 |
| Day 14 | 0.026698 | 26.33 | 21.74 | 4.594 | 1.928 | 2.383 | 21.00 | 0.010786 |
| Day 21 | 0.001300 | 32.11 | 26.32 | 5.787 | 1.560 | 3.709 | 21.00 | 0.000875 |
| Day 28 | 0.001977 | 35.65 | 29.76 | 5.896 | 1.669 | 3.532 | 21.00 | 0.000998 |
| Day 35 | 0.000118 | 40.95 | 31.50 | 9.448 | 1.959 | 4.823 | 19.00 | 0.000239 |
| Day 42 | 0.000296 | 41.74 | 34.88 | 6.864 | 1.467 | 4.681 | 15.00 | 0.000299 |

**Supplementary Table 10:** Both head removal, statistical analysis data.

|  | P value | Mean of Control | Mean of Double decapitation animals | Difference | SE of difference | t ratio | df | q value |
| --- | --- | --- | --- | --- | --- | --- | --- | --- |
| Day 7 | 0.565658 | 19.22 | 18.09 | 1.133 | 1.942 | 0.5832 | 22.00 | 0.380876 |
| Day 14 | 0.018288 | 26.33 | 20.43 | 5.902 | 2.297 | 2.570 | 20.00 | 0.018470 |
| Day 21 | 0.141739 | 32.11 | 28.48 | 3.629 | 2.377 | 1.527 | 21.00 | 0.114525 |
| Day 28 | 0.017421 | 35.65 | 30.76 | 4.892 | 1.887 | 2.592 | 20.00 | 0.018470 |
| Day 35 | 0.000022 | 40.95 | 29.39 | 11.56 | 2.103 | 5.499 | 20.00 | 0.000045 |
| Day 42 | 0.000002 | 41.74 | 28.94 | 12.80 | 1.736 | 7.377 | 15.00 | 0.000009 |

**Supplementary Table 11:**
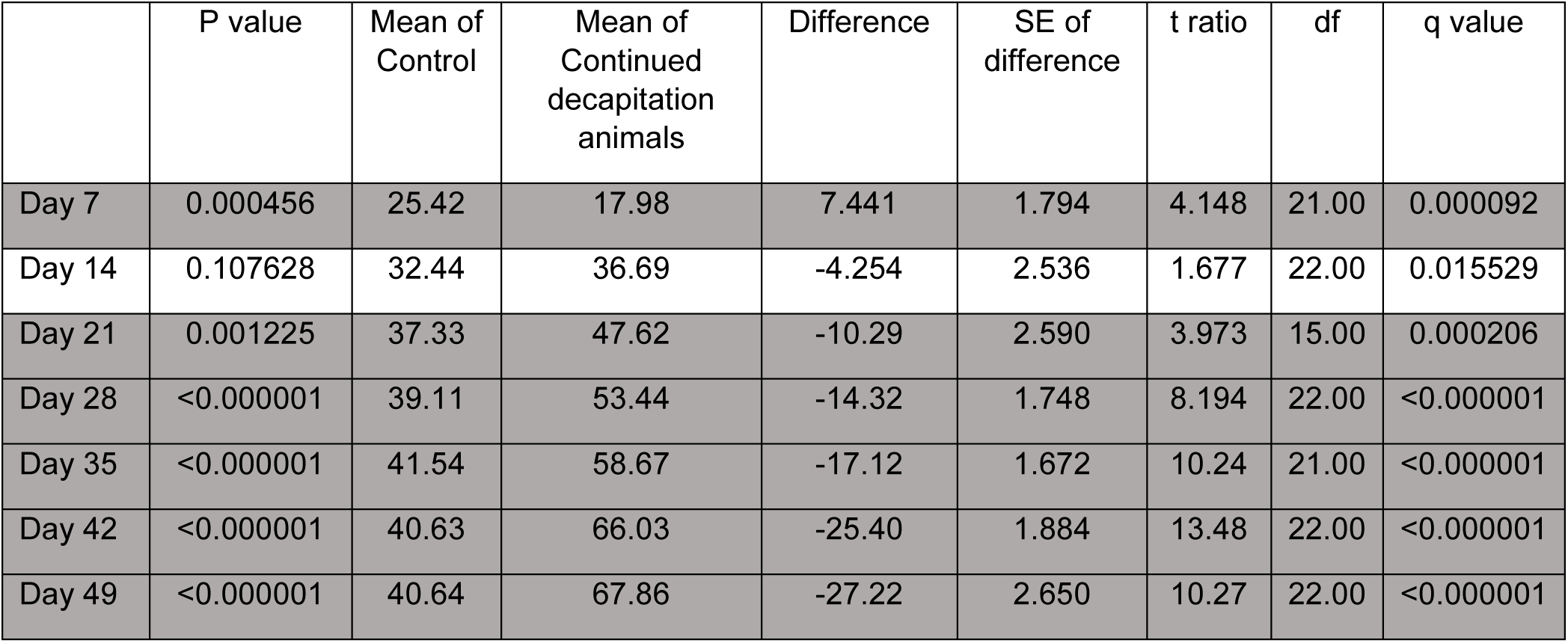
Continuous decapitation, statistical analysis data.

|  | P value | Mean of Control | Mean of Continued decapitation animals | Difference | SE of difference | t ratio | df | q value |
| --- | --- | --- | --- | --- | --- | --- | --- | --- |
| Day 7 | 0.000456 | 25.42 | 17.98 | 7.441 | 1.794 | 4.148 | 21.00 | 0.000092 |
| Day 14 | 0.107628 | 32.44 | 36.69 | -4.254 | 2.536 | 1.677 | 22.00 | 0.015529 |
| Day 21 | 0.001225 | 37.33 | 47.62 | -10.29 | 2.590 | 3.973 | 15.00 | 0.000206 |
| Day 28 | <0.000001 | 39.11 | 53.44 | -14.32 | 1.748 | 8.194 | 22.00 | <0.000001 |
| Day 35 | <0.000001 | 41.54 | 58.67 | -17.12 | 1.672 | 10.24 | 21.00 | <0.000001 |
| Day 42 | <0.000001 | 40.63 | 66.03 | -25.40 | 1.884 | 13.48 | 22.00 | <0.000001 |
| Day 49 | <0.000001 | 40.64 | 67.86 | -27.22 | 2.650 | 10.27 | 22.00 | <0.000001 |

**Supplementary Table 12:** Head tip removal, statistical analysis data.

|  | P value | Mean of Control | Mean of Head tip removal animals | Difference | SE of difference | t ratio | df | q value |
| --- | --- | --- | --- | --- | --- | --- | --- | --- |
| Day 7 | 0.264776 | 18.50 | 20.20 | -1.696 | 1.482 | 1.144 | 22.00 | 0.668559 |
| Day 13 | 0.058891 | 26.71 | 29.83 | -3.121 | 1.566 | 1.992 | 22.00 | 0.297401 |
| Day 24 | 0.749590 | 38.57 | 39.18 | -0.6187 | 1.914 | 0.3232 | 22.00 | 0.900474 |
| Day 28 | 0.711581 | 40.63 | 41.34 | -0.7093 | 1.893 | 0.3748 | 21.00 | 0.900474 |
| Day 35 | 0.891558 | 45.10 | 45.37 | -0.2672 | 1.935 | 0.1381 | 20.00 | 0.900474 |

**Supplementary Table 13:** Mature DH decapitation, statistical analysis data.

| Table Analyzed | DH decapitated at Day 49 |
| --- | --- |
| Column C | Day 70 |
| vs. | vs. |
| Column B | Day 49 |
| **Unpaired t test with Welch's correction** |  |
| P value | 0.0483 |
| P value summary | * |
| Significantly different (P < 0.01)? | No |
| One- or two-tailed P value? | Two-tailed |
| Welch-corrected t, df | t=2.146, df=15.32 |
| **How big is the difference?** |  |
| Mean of column B | 45.89 |
| Mean of column C | 42.24 |
| Difference between means (C - B) ± SEM | -3.650 ± 1.701 |
| 99% confidence interval | -8.646 to 1.346 |
| R squared (eta squared) | 0.2311 |
| **F test to compare variances** |  |
| F, DF<sub>n</sub>, DF<sub>d</sub> | 1.474, 8, 11 |
| P value | 0.5388 |
| P value summary | ns |
| Significantly different (P < 0.01)? | No |
| **Data analyzed** |  |
| Sample size, column B | 12 |
| Sample size, column C | 9 |

**Supplementary Table 14:** Half-head lateral removal, statistical analysis data.

|  | P value | Mean of Control | Mean of HH lateral animals | Difference | SE of difference | t ratio | df | q value |
| --- | --- | --- | --- | --- | --- | --- | --- | --- |
| Day 7 | 0.010864 | 18.09 | 21.84 | -3.752 | 1.349 | 2.782 | 22.00 | 0.002743 |
| Day 14 | 0.000154 | 30.92 | 22.84 | 8.082 | 1.773 | 4.559 | 22.00 | 0.000063 |
| Day 21 | 0.000187 | 36.43 | 29.25 | 7.186 | 1.604 | 4.480 | 22.00 | 0.000063 |
| Day 28 | 0.000108 | 39.00 | 29.22 | 9.774 | 2.057 | 4.752 | 21.00 | 0.000063 |

**Supplementary Table 15:** Half-head removal perpendicular, statistical analysis data.

|  | P value | Mean of Control | Mean of HH perpendicular animals | Difference | SE of difference | t ratio | df | q value |
| --- | --- | --- | --- | --- | --- | --- | --- | --- |
| Day 7 | 0.720203 | 18.50 | 18.99 | -0.4920 | 1.356 | 0.3628 | 22.00 | 0.290962 |
| Day 13 | 0.185634 | 26.71 | 24.50 | 2.208 | 1.616 | 1.366 | 22.00 | 0.093745 |
| Day 24 | 0.000062 | 38.57 | 29.50 | 9.066 | 1.838 | 4.934 | 22.00 | 0.000042 |
| Day 28 | 0.000007 | 40.63 | 30.56 | 10.07 | 1.731 | 5.817 | 22.00 | 0.000008 |
| Day 35 | <0.000001 | 45.10 | 32.18 | 12.92 | 1.359 | 9.508 | 20.00 | <0.000001 |

**Supplementary Table 16:** Relative brain size in irradiated and untreated animals, statistical analysis data.

|  |  |  |
| --- | --- | --- |
| Table Analyzed | Relative brain size | Relative brain size |
| Column M | irradiated Day 28 | irradiated Day 28 |
| vs. | vs. | vs. |
| Column F | Day 21 | Day 30 |
| **Unpaired t test with Welch's correction** |  |  |
| P value | 0.0002 | 0.0005 |
| P value summary | *** | *** |
| Significantly different (P < 0.01)? | Yes | Yes |
| One- or two-tailed P value? | Two-tailed | Two-tailed |
| Welch-corrected t, df | t=4.595, df=19.41 | t=3.919, df=27.10 |
| **How big is the difference?** |  |  |
| Mean of column F | 0.8463 | 0.8412 |
| Mean of column M | 0.6466 | 0.6466 |
| Difference between means (M - F) ± SEM | -0.1997 ± 0.04346 | -0.1945 ± 0.04965 |
| 95% confidence interval | -0.2905 to -0.1089 | -0.2964 to -0.09269 |
| R squared (eta squared) | 0.5210 | 0.3617 |
| **F test to compare variances** |  |  |
| F, DFn, Dfd | 4.407, 17, 58 | 3.503, 17, 12 |
| P value | <0.0001 | 0.0322 |
| P value summary | **** | * |
| Significantly different (P < 0.05)? | Yes | Yes |
| **Data analyzed** |  |  |
| Sample size, column F | 59 | 13 |
| Sample size, column M | 18 | 18 |

**Supplementary Table 17:** Head bisection, statistical analysis data.

|  | P value | Mean of Control | Mean of Head bisection animals | Difference | SE of difference | t ratio | df | q value |
| --- | --- | --- | --- | --- | --- | --- | --- | --- |
| Day 7 | 0.666008 | 18.50 | 19.05 | -0.5484 | 1.253 | 0.4375 | 22.00 | 0.134534 |
| Day 13 | 0.002632 | 26.71 | 20.05 | 6.662 | 1.965 | 3.390 | 22.00 | 0.000665 |
| Day 24 | 0.000001 | 38.57 | 28.39 | 10.17 | 1.559 | 6.523 | 22.00 | <0.000001 |
| Day 28 | <0.000001 | 40.63 | 27.33 | 13.30 | 1.801 | 7.382 | 22.00 | <0.000001 |
| Day 35 | <0.000001 | 45.10 | 28.46 | 16.64 | 1.900 | 8.760 | 19.00 | <0.000001 |

**Supplementary Table 18:** Lateral cut at primary head, statistical analysis data.

|  | P value | Mean of Control | Mean of cut at 1st head animals | Difference | SE of difference | t ratio | df | q value |
| --- | --- | --- | --- | --- | --- | --- | --- | --- |
| Day 7 | 0.143374 | 19.76 | 17.25 | 2.512 | 1.655 | 1.518 | 22.00 | 0.086885 |
| Day 14 | 0.002353 | 27.34 | 30.84 | -3.503 | 1.088 | 3.220 | 46.00 | 0.005497 |
| Day 21 | 0.007365 | 33.29 | 36.72 | -3.435 | 1.224 | 2.807 | 45.00 | 0.007439 |
| Day 28 | 0.003628 | 39.22 | 43.97 | -4.755 | 1.543 | 3.081 | 42.00 | 0.005497 |
| Day 35 | 0.141438 | 42.56 | 45.67 | -3.109 | 2.003 | 1.552 | 15.00 | 0.086885 |

**Supplementary Table 19:** Lateral cut at secondary head, statistical analysis data.

|  | P value | Mean of Control | Mean of cut at 2nd head animals | Difference | SE of difference | t ratio | df | q value |
| --- | --- | --- | --- | --- | --- | --- | --- | --- |
| Day 7 | 0.894110 | 19.76 | 19.96 | -0.1972 | 1.465 | 0.1347 | 22.00 | 0.361220 |
| Day 14 | 0.018651 | 27.34 | 24.60 | 2.738 | 1.121 | 2.443 | 44.00 | 0.009419 |
| Day 21 | 0.000153 | 33.29 | 28.33 | 4.951 | 1.200 | 4.127 | 46.00 | 0.000154 |
| Day 28 | <0.000001 | 39.22 | 29.24 | 9.974 | 1.351 | 7.385 | 46.00 | <0.000001 |
| Day 35 | 0.000335 | 42.56 | 30.81 | 11.75 | 2.663 | 4.414 | 18.00 | 0.000226 |

**Supplementary Table 20:**
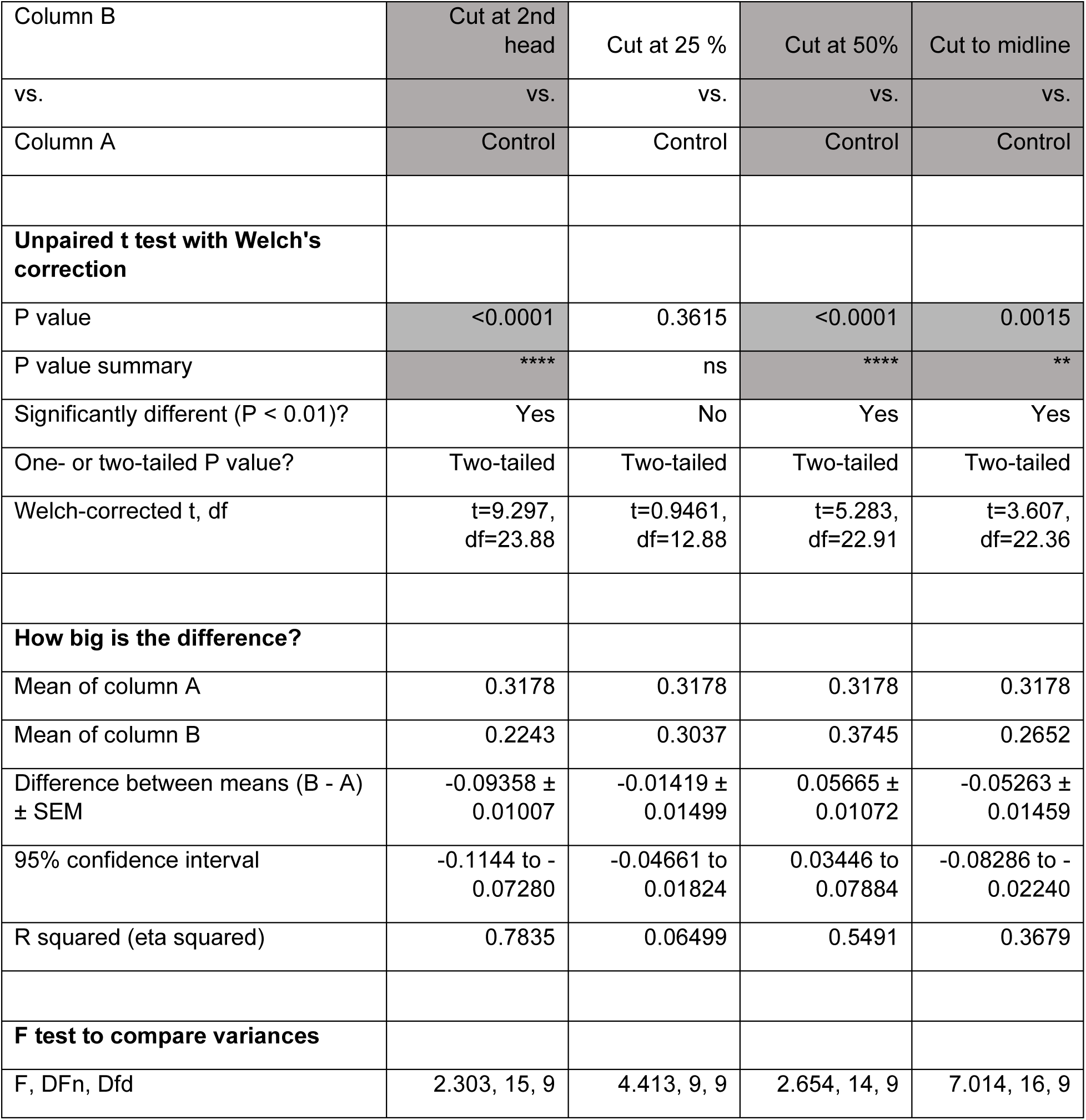
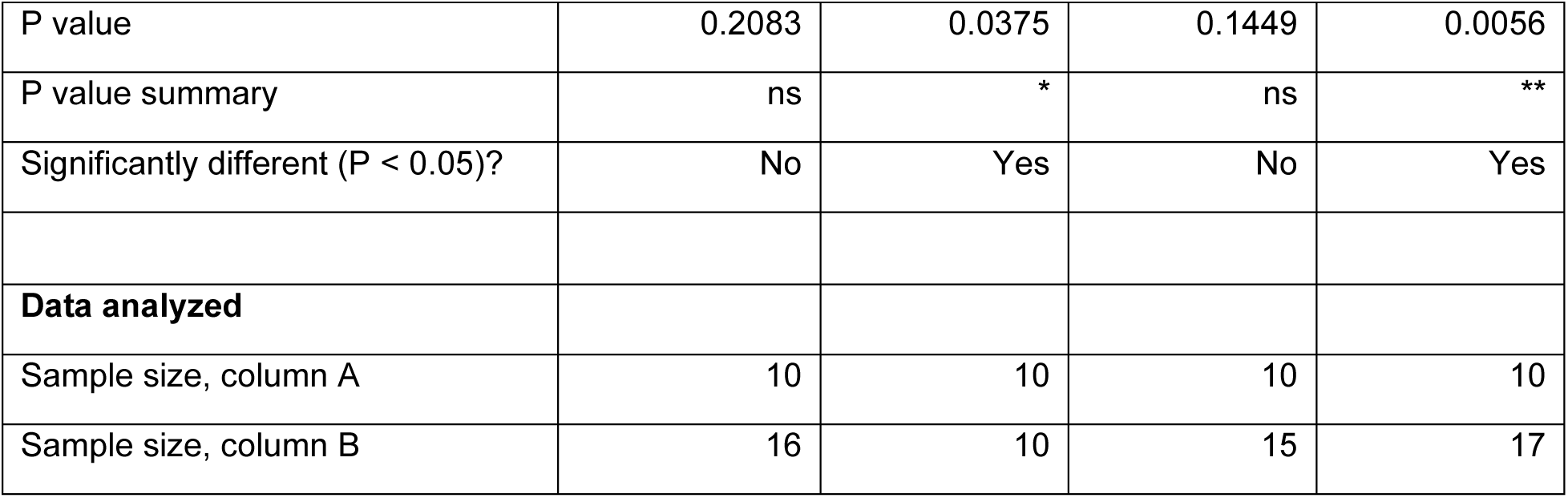
Lateral cut at different positions along the body axis, statistical analysis data.

| Column B | Cut at 2nd head | Cut at 25 % | Cut at 50% | Cut to midline |
| --- | --- | --- | --- | --- |
| vs. | vs. | vs. | vs. | vs. |
| Column A | Control | Control | Control | Control |
| **Unpaired t test with Welch's correction** |  |  |  |  |
| P value | <0.0001 | 0.3615 | <0.0001 | 0.0015 |
| P value summary | **** | ns | **** | ** |
| Significantly different (P < 0.01)? | Yes | No | Yes | Yes |
| One- or two-tailed P value? | Two-tailed | Two-tailed | Two-tailed | Two-tailed |
| Welch-corrected t, df | t=9.297, df=23.88 | t=0.9461, df=12.88 | t=5.283, df=22.91 | t=3.607, df=22.36 |
| **How big is the difference?** |  |  |  |  |
| Mean of column A | 0.3178 | 0.3178 | 0.3178 | 0.3178 |
| Mean of column B | 0.2243 | 0.3037 | 0.3745 | 0.2652 |
| Difference between means (B - A) ± SEM | -0.09358 ± 0.01007 | -0.01419 ± 0.01499 | 0.05665 ± 0.01072 | -0.05263 ± 0.01459 |
| 95% confidence interval | -0.1144 to -0.07280 | -0.04661 to 0.01824 | 0.03446 to 0.07884 | -0.08286 to -0.02240 |
| R squared (eta squared) | 0.7835 | 0.06499 | 0.5491 | 0.3679 |
| **F test to compare variances** |  |  |  |  |
| F, DFn, Dfd | 2.303, 15, 9 | 4.413, 9, 9 | 2.654, 14, 9 | 7.014, 16, 9 |
| P value | 0.2083 | 0.0375 | 0.1449 | 0.0056 |
| P value summary | ns | * | ns | ** |
| Significantly different (P < 0.05)? | No | Yes | No | Yes |
| **Data analyzed** |  |  |  |  |
| Sample size, column A | 10 | 10 | 10 | 10 |
| Sample size, column B | 16 | 10 | 15 | 17 |

**Supplementary Table 21:** Regenerative outcomes in DHs of different ages and cutting planes, statistical analysis data.

|  | P value | Mean of new DH | Mean of old DH | Difference | SE of difference | t ratio | df | q value |
| --- | --- | --- | --- | --- | --- | --- | --- | --- |
| 2nd decapitation | 0.744677 | 95.50 | 98.25 | -2.750 | 8.372 | 0.3285 | 32.00 | 0.300850 |
| 1/4 body length cut - primary side | <0.000001 | 31.75 | 86.75 | -55.00 | 8.372 | 6.570 | 32.00 | <0.000001 |
| 1/4 body length cut - secondary side | 0.002898 | 29.00 | 2.000 | 27.00 | 8.372 | 3.225 | 32.00 | 0.001951 |
| midway cut - primary half | 0.122455 | 3.400 | 16.00 | -12.60 | 7.942 | 1.587 | 32.00 | 0.061840 |
| midway cut - secondary half | <0.000001 | 96.80 | 16.00 | 80.80 | 7.942 | 10.17 | 32.00 | <0.000001 |

## References

Adell, T., Cebria, F., & Salo, E. (2010). Gradients in planarian regeneration and homeostasis. Cold Spring Harb Perspect Biol, 2(1), a000505. doi:10.1101/cshperspect.a000505

Adler, C. E., Seidel, C. W., McKinney, S. A., & Sanchez Alvarado, A. (2014). Selective amputation of the pharynx identifies a FoxA-dependent regeneration program in planaria. Elife, 3, e02238. doi:10.7554/eLife.02238

Azimzadeh, J., & Basquin, C. (2016). Basal bodies across eukaryotes series: basal bodies in the freshwater planarian Schmidtea mediterranea. Cilia, 5, 15. doi:10.1186/s13630-016-0037-1

Azimzadeh, J., Wong, M. L., Downhour, D. M., Sanchez Alvarado, A., & Marshall, W. F. (2012). Centrosome loss in the evolution of planarians. Science, 335(6067), 461–463. doi:10.1126/science.1214457

Boisvieux-Ulrich, E., Sandoz, D., & Allart, J.-P. (1991). Determination of ciliary polarity precedes differentiation in the epithelial cells of quail oviduct. Biology of the Cell, 72(1-2), 3–14.

Brooks, E. R., & Wallingford, J. B. (2014). Multiciliated cells. Curr Biol, 24(19), R973–982. doi:10.1016/j.cub.2014.08.047

Chien, Y.-H., Werner, M. E., Stubbs, J., Joens, M. S., Li, J., Chien, S., … Kintner, C. (2013). Bbof1 is required to maintain cilia orientation. Development, 140(16), 3468. doi:10.1242/dev.096727

Chien, Y. H., Srinivasan, S., Keller, R., & Kintner, C. (2018). Mechanical Strain Determines Cilia Length, Motility, and Planar Position in the Left-Right Organizer. Dev Cell, 45(3), 316–330 e314. doi:10.1016/j.devcel.2018.04.007

Currie, K. W., & Pearson, B. J. (2013). Transcription factors lhx1/5-1 and pitx are required for the maintenance and regeneration of serotonergic neurons in planarians. Development, 140(17), 3577–3588. doi:10.1242/dev.098590

Davey, C. F., & Moens, C. B. (2017). Planar cell polarity in moving cells: think globally, act locally. Development, 144(2), 187. doi:10.1242/dev.122804

Delhaas, T., Decaluwe, W., Rubbens, M., Kerckhoffs, R., & Arts, T. (2004). Cardiac fiber orientation and the left-right asymmetry determining mechanism. Annals of the New York Academy of Sciences, 1015, 190–201.

Devenport, D. (2014). The cell biology of planar cell polarity. The Journal of Cell Biology, 207(2), 171. doi:10.1083/jcb.201408039

Durant, F., Morokuma, J., Fields, C., Williams, K., Adams, D. S., & Levin, M. (2017). Long-Term, Stochastic Editing of Regenerative Anatomy via Targeting Endogenous Bioelectric Gradients. Biophys J, 112(10), 2231–2243. doi:10.1016/j.bpj.2017.04.011

Fields, C., Bischof, J., & Levin, M. (2019). Morphological coordination: A common ancestral function unifying neural and non-neural signaling. Physiology, in print.

Goodrich, L. V. (2008). The plane facts of PCP in the CNS. Neuron, 60(1), 9–16. doi:10.1016/j.neuron.2008.09.003

Herrera-Rincon, C., Pai, V. P., Moran, K. M., Lemire, J. M., & Levin, M. (2017). The brain is required for normal muscle and nerve patterning during early Xenopus development. Nat Commun, 8(1), 587. doi:10.1038/s41467-017-00597-2

Iwadate, Y., & Suzaki, T. (2004). Ciliary reorientation is evoked by a rise in calcium level over the entire cilium. Cell Motility and the Cytoskeleton, 57(4), 197–206. doi:10.1002/cm.10165

King, S. M., & Patel-King, R. S. (2016). Planaria as a Model System for the Analysis of Ciliary Assembly and Motility. Methods Mol Biol, 1454, 245–254. doi:10.1007/978-1-4939-3789-9_16

Klagges, B. R. E., Heimbeck, G., Godenschwege, T. A., Hofbauer, A., Pflugfelder, G. O., Reifegerste, R., … Buchner, E. (1996). Invertebrate Synapsins: A Single Gene Codes for Several Isoforms in Drosophila. The Journal of Neuroscience, 16(10), 3154–3165. doi:10.1523/jneurosci.16-10-03154.1996

Konishi, S., Gotoh, S., Tateishi, K., Yamamoto, Y., Korogi, Y., Nagasaki, T., … Mishima, M. (2016). Directed Induction of Functional Multi-ciliated Cells in Proximal Airway Epithelial Spheroids from Human Pluripotent Stem Cells. Stem Cell Reports, 6(1), 18–25. doi:10.1016/j.stemcr.2015.11.010

Kunimoto, K., Yamazaki, Y., Nishida, T., Shinohara, K., Ishikawa, H., Hasegawa, T., … Tsukita, S. (2012). Coordinated ciliary beating requires Odf2-mediated polarization of basal bodies via basal feet. Cell, 148(1-2), 189–200. doi:10.1016/j.cell.2011.10.052

Lee, M., & Vasioukhin, V. (2008). Cell polarity and cancer--cell and tissue polarity as a non-canonical tumor suppressor. J Cell Sci, 121(Pt 8), 1141–1150. doi:10.1242/jcs.016634

Marshall, W. F., & Kintner, C. (2008). Cilia orientation and the fluid mechanics of development. Curr Opin Cell Biol, 20(1), 48–52. doi:10.1016/j.ceb.2007.11.009

Marz, M., Seebeck, F., & Bartscherer, K. (2013). A Pitx transcription factor controls the establishment and maintenance of the serotonergic lineage in planarians. Development, 140(22), 4499–4509. doi:10.1242/dev.100081

Meunier, A., & Azimzadeh, J. (2016). Multiciliated Cells in Animals. Cold Spring Harbor perspectives in biology, 8(12), a028233. doi:10.1101/cshperspect.a028233

Mitchell, B., Jacobs, R., Li, J., Chien, S., & Kintner, C. (2007). A positive feedback mechanism governs the polarity and motion of motile cilia. Nature, 447(7140), 97–101. doi:10.1038/nature05771

Morgan, T. H. (1901). Regeneration. New York: Macmillan.

Noguchi, M., Kitani, T., Ogawa, T., Inoue, H., & Kamachi, H. (2005). Augmented ciliary reorientation response and cAMP-dependent protein phosphorylation induced by glycerol in triton-extracted Paramecium. Zoolog Sci, 22(1), 41–48. doi:10.2108/zsj.22.41

Ohata, S., & Alvarez-Buylla, A. (2016). Planar Organization of Multiciliated Ependymal (E1) Cells in the Brain Ventricular Epithelium. Trends Neurosci, 39(8), 543–551. doi:10.1016/j.tins.2016.05.004

Oviedo, N. J., Morokuma, J., Walentek, P., Kema, I. P., Gu, M. B., Ahn, J. M., … Levin, M. (2010). Long-range neural and gap junction protein-mediated cues control polarity during planarian regeneration. Dev Biol, 339(1), 188–199. doi:10.1016/j.ydbio.2009.12.012

Oviedo, N. J., Nicolas, C. L., Adams, D. S., & Levin, M. (2008). Establishing and maintaining a colony of planarians. CSH Protoc, 2008, pdb prot5053. doi:10.1101/pdb.prot5053

Pan, J., You, Y., Huang, T., & Brody, S. L. (2007). RhoA-mediated apical actin enrichment is required for ciliogenesis and promoted by Foxj1. J Cell Sci, 120(Pt 11), 1868–1876. doi:10.1242/jcs.005306

Panizzi, J. R., Jessen, J. R., Drummond, I. A., & Solnica-Krezel, L. (2007). New functions for a vertebrate Rho guanine nucleotide exchange factor in ciliated epithelia. Development, 134(5), 921. doi:10.1242/dev.02776

Pearl, R. (1903). The movements and reactions of fresh-water planarians: a study in animal behaviour. Journal of Cell Science, 46.

Petersen, C. P., & Reddien, P. W. (2008). Smed-betacatenin-1 is required for anteroposterior blastema polarity in planarian regeneration. Science, 319(5861), 327–330. doi:10.1126/science.1149943

Petersen, C. P., & Reddien, P. W. (2011). Polarized activation of notum at wounds inhibits Wnt Function to Promote Planarian Head Regeneration. Science, 332.

Pietak, A., Bischof, J., LaPalme, J., Morokuma, J., & Levin, M. (2019). Neural control of body-plan axis in regenerating planaria. PLoS Comput Biol, 15(4), e1006904. doi:10.1371/journal.pcbi.1006904

Reddien, P. W. (2018). The Cellular and Molecular Basis for Planarian Regeneration. Cell, 175(2), 327–345. doi:10.1016/j.cell.2018.09.021

Reddien, P. W., & Sanchez Alvarado, A. (2004). Fundamentals of planarian regeneration. Annu Rev Cell Dev Biol, 20, 725–757. doi:10.1146/annurev.cellbio.20.010403.095114

Rompolas, P., Azimzadeh, J., Marshall, W. F., & King, S. M. (2013). Analysis of ciliary assembly and function in planaria. Methods Enzymol, 525, 245–264. doi:10.1016/B978-0-12-397944-5.00012-2

Rompolas, P., Patel-King, R. S., & King, S. M. (2010). An outer arm Dynein conformational switch is required for metachronal synchrony of motile cilia in planaria. Mol Biol Cell, 21(21), 3669–3679. doi:10.1091/mbc.E10-04-0373

Rueden, C. T., Schindelin, J., Hiner, M. C., DeZonia, B. E., Walter, A. E., Arena, E. T., & Eliceiri, K. W. (2017). ImageJ2: ImageJ for the next generation of scientific image data. BMC Bioinformatics, 18(1), 529. doi:10.1186/s12859-017-1934-z

Rustia, C. P. (1925). The control of biaxial development in the reconstitution of pieces of Planaria. Journal of Experimental Zoology, 42(1), 111–142. doi:DOI 10.1002/jez.1400420106

Schindelin, J., Arganda-Carreras, I., Frise, E., Kaynig, V., Longair, M., Pietzsch, T., … Cardona, A. (2012). Fiji: an open-source platform for biological-image analysis. Nat Methods, 9(7), 676–682. doi:10.1038/nmeth.2019

Schliwa, M. (1982). Action of cytochalasin D on cytoskeletal networks. The Journal of Cell Biology, 92, 79–91.

Spassky, N., & Meunier, A. (2017). The development and functions of multiciliated epithelia. Nature Reviews Molecular Cell Biology, 18, 423. doi:10.1038/nrm.2017.21

Stevenson, C. G., & Beane, W. S. (2010). A low percent ethanol method for immobilizing planarians. PLoS One, 5(12), e15310. doi:10.1371/journal.pone.0015310

Thielicke, W., & Stamhuis, E. J. (2014). PIVlab – Towards User-friendly, Affordable and Accurate Digital Particle Image Velocimetry in MATLAB. Journal of Open Research Software, 2. doi:10.5334/jors.bl

Thielicke, W., & Stamhuis, E. J. (2019). PIVlab - Time-Resolved Digital Particle Image Velocimetry Tool for MATLAB.

Tissir, F., & Goffinet, A. M. (2010). Planar cell polarity signaling in neural development. Curr Opin Neurobiol, 20(5), 572–577. doi:10.1016/j.conb.2010.05.006

Tung, T.-C., & Yeh-Tung, Y. (1940). Experimental studies on the determination of polarity of ciliary action of anuran embryos: H. Vaillant-Carmanne.

Twitty. (1928). Experimental studies on the ciliary action of amphibian embryos. Journal of Experimental Zoology, 50(3), 319–344.

Veraszto, C., Ueda, N., Bezares-Calderon, L. A., Panzera, A., Williams, E. A., Shahidi, R., & Jekely, G. (2017). Ciliomotor circuitry underlying whole-body coordination of ciliary activity in the Platynereis larva. Elife, 6. doi:10.7554/eLife.26000

Vu, H. T.-K., Mansour, S., Kuecken, M., Blasse, C., Basquin, C., Azimzadeh, J., … Rink, J. C. (2018). Multi-scale coordination of planar cell polarity in planarians. bioRxiv. doi:10.1101/324822

Wagner, D. E., Wang, I. E., & Reddien, P. W. (2011). Clonogenic Neoblasts Are Pluripotent Adult Stem Cells That Underlie Planarian Regeneration. Science, 332(6031), 811–816. doi:10.1126/science.1203983

Wallingford, J. B. (2010). Planar cell polarity signaling, cilia and polarized ciliary beating. Curr Opin Cell Biol, 22(5), 597–604. doi:10.1016/j.ceb.2010.07.011

Wallingford, J. B. (2012). Planar cell polarity and the developmental control of cell behavior in vertebrate embryos. Annu Rev Cell Dev Biol, 28, 627–653. doi:10.1146/annurev-cellbio-092910-154208

Werner, M. E., Hwang, P., Huisman, F., Taborek, P., Yu, C. C., & Mitchell, B. J. (2011). Actin and microtubules drive differential aspects of planar cell polarity in multiciliated cells. J Cell Biol, 195(1), 19–26. doi:10.1083/jcb.201106110

Werner, M. E., & Mitchell, B. J. (2012a). Planar cell polarity: microtubules make the connection with cilia. Curr Biol, 22(23), R1001–1004. doi:10.1016/j.cub.2012.10.030

Werner, M. E., & Mitchell, B. J. (2012b). Understanding ciliated epithelia: the power of Xenopus. Genesis, 50(3), 176–185. doi:10.1002/dvg.20824

Witchley, J. N., Mayer, M., Wagner, D. E., Owen, J. H., & Reddien, P. W. (2013). Muscle cells provide instructions for planarian regeneration. Cell Rep, 4(4), 633–641. doi:10.1016/j.celrep.2013.07.022

